# CD226^+^ macrophages arise from MDP-derived monocytes and regulate lipid metabolism

**DOI:** 10.1101/2024.12.03.626330

**Authors:** Alexandre Gallerand, Zakariya Caillot, Frederike Westermann, Selma Tuzlak, Sacha Grenet, Marina Terekhova, Alexia Castiglione, Thomas Pilot, Laurent L’homme, Kerim Acil, Lorlana Leporati, Margaux Giacchero, Eloïse Goës, Maxime Franceschini, Sebastien Fleury, Evy Boré, Michael Chang, Bastien Dolfi, Fairouz N. Zair, Audrey Bennetot, Zandile Mlamla, David Voehringer, Matthias Mack, Jaap G. Neels, Joel Haas, David Dombrowicz, Jesse W. Williams, David Masson, Elvira Mass, Maxim N. Artyomov, Burkhard Becher, Adeline Bertola, Stoyan Ivanov

## Abstract

Macrophages are innate immune cells present in all tissues, in which they participate in immune responses and maintenance of tissue homeostasis. They develop either from embryonic precursors or from circulating monocytes, and their origin impacts their functions. We previously observed robust recruitment of monocytes to brown adipose tissue in which they could differentiation into two distinct macrophage subsets identifiable by CD206 or CD226 expression. In the present study, we investigated monocyte differentiation pathways in brown adipose tissue and the function of monocyte-derived macrophages. Fate mapping analysis revealed a low contribution of GMP- and a high contribution of MDP-derived monocytes to the CD226^high^ macrophage subset. Importantly, adoptive transfer experiments demonstrate that MDP- but not GMP-derived monocytes are pre-conditioned to give rise to CD226^high^ macrophages. We found that MDP-derived CD226^high^ macrophages were also present in other tissues including peritoneal cavity, adrenal glands and all adipose depots. CD226^high^ macrophages were regulated by both GM-CSF and CSF1R. Genetic depletion of CD226^high^ macrophages caused increased BAT and plasma triglyceride content. We thus identify CD226^high^ MDP-derived macrophages as a new myeloid cell type conserved across tissues and tied to lipid metabolism homeostasis.

## Introduction

Macrophages are innate immune cells observed in all organs ^1^. These cells originate from embryonic or bone marrow precursors ^1,2^. In adulthood, the proportion of monocyte-and embryonically-derived macrophages varies in different tissues. For example, brain macrophages are exclusively derived from embryonic precursors and their pool is sustained through proliferation, while in gut, lung and spleen, macrophages are more frequently renewed by recruitment of blood monocytes ^2,3^. While brown adipose tissue macrophages are initially seeded during embryogenesis ^4^, they are diluted through monocyte recruitment after birth ^4,5^.

Brown adipose tissue (BAT) is actively involved in heat generation by adaptive thermogenesis ^6^. UCP1 (Uncoupling Protein 1) expression and high mitochondria content are BAT hallmarks ^7^. Previous work established that BAT contains several macrophage subsets that heavily rely on CCR2-dependent monocyte recruitment for their maintenance ^5^. Nevertheless, the functions of BAT macrophage subsets remain ill-characterized. A subset of CX3CR1^+^ BAT macrophages regulates neuron network density and subsequently the thermogenic response ^8^. Adipose tissue macrophage depletion with a CSF1R-blocking antibody was associated with a decreased local lipid content ^9^. Yet, whether this function is fulfilled by a selective macrophage subset, or requires the coordinated contribution of several macrophage populations, is not yet established.

Monocytes are generated in the bone marrow through a tight spatially and temporally regulated multistep process named myelopoiesis. Two major monocyte subsets are present in mice and express distinct phenotypic markers ^10,11^. Ly6C^high^ classical monocytes are recruited to peripheral tissues at steady state and upon inflammation and contribute to the maintenance of tissue resident macrophages ^3^. Ly6C^low^ non-classical monocytes contribute to blood vessel health by secreting growth factors and play a pivotal role during cancer metastasis ^12–14^. The majority of blood monocytes originate from granulocyte-monocyte precursors (GMPs) ^3^. However, an alternative pathway generating monocytes and dependent on monocyte-dendritic cell precursors (MDPs) has been also proposed ^15,16^. We previously established that BAT macrophages are constantly supplied by blood monocytes, yet whether GMP- and MDP-derived monocytes give rise preferentially or exclusively to selective BAT macrophage subsets is yet not established.

In the present study, we report that CD226^high^ macrophages are a unique macrophage subset that is conserved across tissues, including adipose tissues and serosal cavities. These cells particularly accumulate in thermogenic adipose depots in which they regulate triglyceride homeostasis. CD226^high^ macrophages arise in majority from MDP-derived monocytes. Invalidating *Csf1r* or *Csf2* leads to a partial reduction of CD226^high^ macrophage numbers in adipose tissue and peritoneal cavity, suggesting that this atypical macrophage population relies on both CSF1 and CSF2 for their generation and maintenance. Together, these results indicate that the CD226^high^ macrophages are a unique myeloid cell population controlling lipid metabolism both locally and systemically.

## Results

### CD226^high^ macrophages arise from monocytes but show low contribution from GMPs

We recently demonstrated that BAT contained numerous immune cells by using single-cell RNA sequencing (scRNA-seq) analysis ^5^. BAT contained several monocyte, macrophage and dendritic cell (DC) clusters (**Figure S1a**). Among those cells, *Itgam* (CD11b) mRNA was expressed by the majority of the clusters (**Figure S1b**). We identified three monocyte subsets (clusters 2, 3 and 4) in BAT characterized by their relative expression of *Ly6c2* and *Nr4a1* mRNA (**Figure S1b**). Cluster 2 contained Ly6C^low^ monocytes and cluster 4 included Ly6C^high^ monocytes. Furthermore, we predicted that BAT contained 4 well-defined macrophage subsets (clusters 5, 6, 7 and 8). Cluster 5 macrophages expressed *Cd226* mRNA and several genes encoding for proteins involved in extracellular matrix remodeling and these cells were initially named “Matrix” macrophages (**Figure S1c**) ^5^. Cluster 6 was the larger macrophage cluster (**Figure S1c**). Cluster 6 macrophages highly expressed genes previously associated with macrophage alternative polarization in response to IL-4 (*Mrc1*, *Lyve1*, *CD163*) (**Figure S1c**). These cells were called CD206^high^ macrophages (**Figure S1c**) ^5^. Two additional macrophage clusters were also identified (clusters 7 and 8) and were characterized by the expression of enzymes involved in lipid metabolism (*Lpl* and *Plin2*). To validate at the protein level the predicted myeloid cell diversity in BAT, we developed a gating strategy and used spectral flow cytometry. Macrophages were identified as CD45^+^ F4/80^+^ MerTK^+^ cells (**Figure S1d**). According to our scRNA-seq data predictions, matrix macrophages were found to highly express CD226 and had low CD64 (Fcgr1) expression (**Figures 1a, S1c-d**). CD206^high^ macrophages were found to highly express both CD206 and CD64 (**Figures S1c-e**). A fraction of these cells also co-expressed Lyve1 and Timd4, suggesting long tissue residency (**Figures S1c-e**). Uptake of intravenously injected TRITC-Dextran indicated that CD206^high^ macrophages displayed phagocytic behavior and were likely associated with blood vessels while CD226^high^ macrophages showed lower dextran uptake (**Figure S1f**). Since we could not define distinct membrane markers for macrophage clusters 7 and 8, these cells were identified as CD45^+^ F4/80^+^ MerTK^+^ CD206^-^ CD226^-^ cells (DN macrophages). Ly6C^high^ monocytes were defined as CD45^+^ MerTK^-^ Ly6C^+^ CD11b^+^ Ly6G^-^ cells (**Figure S1d**). We also found cells corresponding to conventional DC (cDC) subsets. cDC2 were identified as CD45^+^ MerTK^-^ CD11c^+^ MHC-II^+^ CD11b^+^ cells while cDC1 were CD45^+^ MerTK^-^ CD11c^+^ MHC-II^+^ CD24^+^ (**Figure S1d**).

**Figure 1.**
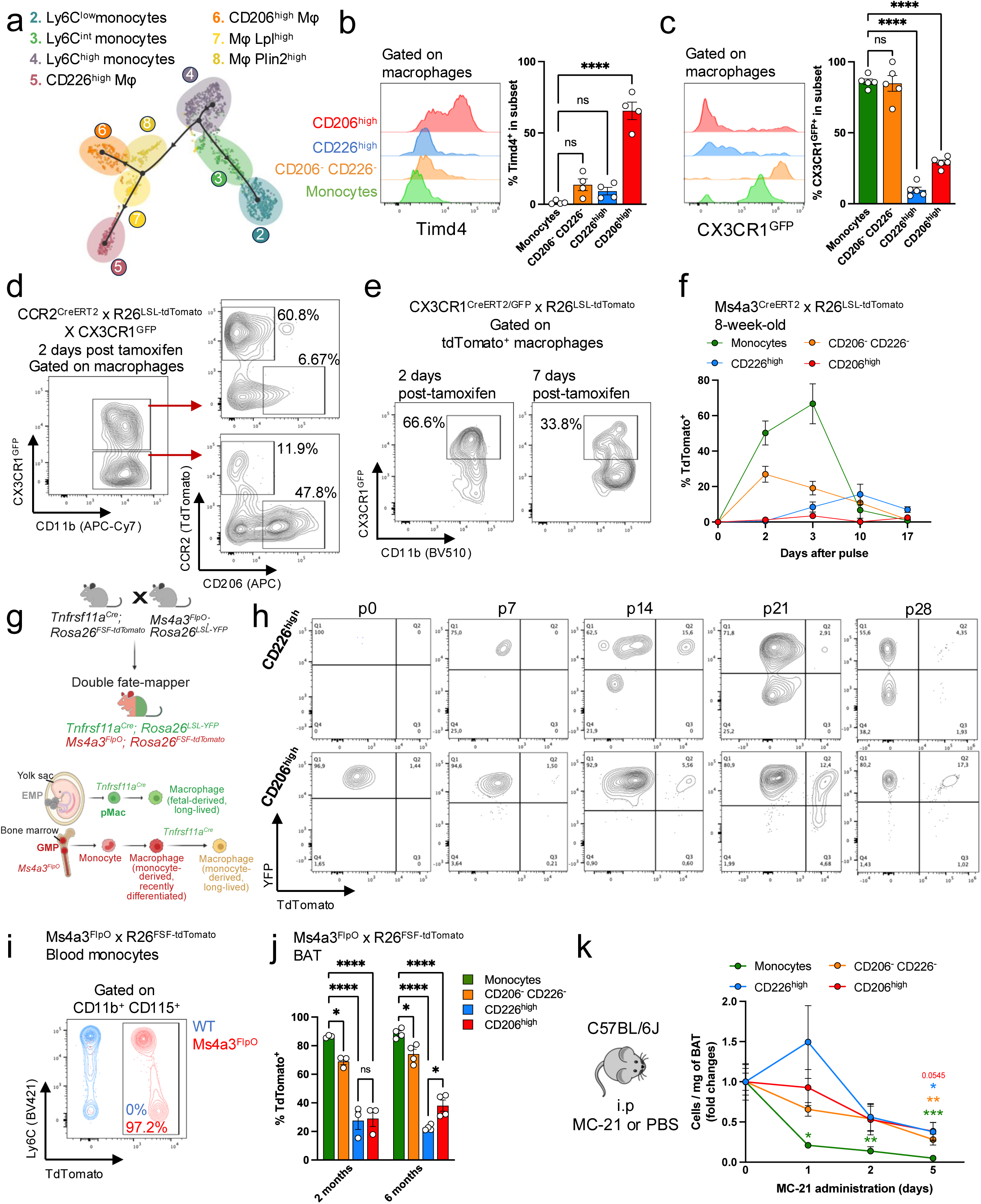
Characterization of brown adipose tissue CD226^high^ macrophages. (**a**) Trajectory analysis of BAT myeloid cells using Slingshot. (**b, c**) Analysis of Timd4 (b) and CX3CR1^GFP^ (c) expression in BAT macrophages. (**d**) Fate mapping analysis of monocyte contribution to CX3CR1^+^ and CX3CR1^-^ BAT macrophages in CCR2^CreERT2^; R26^tdTomato^; CX3CR1^GFP^ mice that received tamoxifen gavage two days prior. (**e**) Fate mapping analysis of CX3CR1^+^ BAT macrophages using double reporter CX3CR1^CreERT2/GFP^; R26^tdTomato^ mice. (**f**) Time-course analysis of BAT monocyte and macrophage tdTomato labeling following tamoxifen administration in Ms4a3^CreERT2^; R26^tdTomato^ mice. (**g**) Schematic representation of the macrophage lineage tracing strategy using the double fate-mapping model Tnfrsf11a^Cre^; Rosa26^LSL-YFP^; Ms4a3^FlpO^; Rosa26^FSF-tdTomato^. *Tnfrsf11a* is expressed in pre-macrophages (pM ac) originating from yolk sac erythro-myeloid progenitors (EMP). Long-lived macrophages derived from the yolk sac are marked by the yellow fluorescent protein (YFP) signal, depicted in green. *Ms4a3* is expressed in granulocyte-monocyte progenitors (GMP), which arise from hematopoietic stem cells (HSC) during definitive hematopoiesis. Short-lived or recently differentiated monocyte-derived macrophages are labeled with tdTomato, represented in red. If a monocyte-derived macrophage persists in the tissue, *Tnfrsf11a* is upregulated, triggering recombination at the Rosa26^LSL-YFP^ locus and resulting in co-expression of both YFP and tdTomato, indicated by a yellow signal. (**h**) Flow cytometry analysis of tdTomato and YFP expression in BAT CD226^high^ and CD206^high^ macrophages from Ms4a3^Flp^; R26^FSF-tdTomato^; Tnfrsf11a^Cre^; R26^LSL-YFP^ mice. (**i**) Flow cytometry analysis of tdTomato expression in blood monocytes from Ms4a3^Flp^; R26^FSF-tdTomato^ mice. (**j**) Quantification of tdTomato^+^ cells among the indicated BAT monocyte and macrophage populations in Ms4a3^Flp^; R26^FSF-tdTomato^ mice. (**k**) Time-course quantification of BAT macrophage subsets along MC-21 administration. Data are presented as mean values +/- SEM. See also Figures S1 and S2.

Next, we aimed at defining the developmental connections between macrophage subsets. We previously demonstrated a high contribution of blood monocytes to the BAT macrophage pool ^5^. Using Slingshot analysis of BAT monocyte and macrophage clusters, we predicted that monocytes could generate two terminally differentiated BAT populations: CD226^high^ and CD206^high^ macrophages (**Figure 1a**). Macrophage clusters 7 and 8 appeared as intermediate stages of differentiation towards the generation of clusters 5 and 6 (**Figure 1a**). Importantly, Timd4 expression was high selectively in CD206^high^ macrophages, suggesting that these cells could be, at least partially, from an embryonic origin (**Figure 1b**). Timd4 expression was low on DN and CD226^high^ macrophages further supporting their monocyte origin (**Figure 1b**). Our scRNA-seq data revealed that *Cx3cr1* mRNA was highly expressed in BAT monocytes but not macrophages (**Figure S1c**). To assess CX3CR1 protein expression on BAT myeloid cells, we used CX3CR1^GFP/+^ mice ^17^. BAT monocytes and DN macrophages highly expressed CX3CR1 (**Figure 1c**). A fraction of CD206^high^ macrophages (nearly 25%) was also GFP^+^ (**Figure 1c**). GFP expression level was the lowest in BAT CD226^high^ macrophages (**Figure 1c**). These observations suggested that monocytes could differentially contribute to BAT macrophage subsets, with a lower contribution to the CD206^high^ macrophage subset.

We next used genetic pulse-chase experiments to investigate monocyte contribution to BAT macrophage populations. For this purpose, *CCR2^CreERT^*^2^*; R26^LSL-tdT^; CX3CR1^GFP/+^* mice were administered with tamoxifen (TAM) and analyzed 48 hours later (**Figure 1d**). We analyzed the appearance of tdTomato^+^ cells in macrophage subsets and found that around 60% of CX3CR1^GFP+^ cells were labelled at this time point (**Figure 1d**). Importantly, the majority of CX3CR1-expressing CD206^high^ BAT macrophages were not tdTomato^+^, suggesting that this population is not monocyte-derived, or its replenishment rate is slower in comparison to CD206^-^ macrophages (**Figure 1d**). Among CX3CR1^-^ BAT macrophages, tdTomato labelling was lower, as only around 12% were marked, and these cells were CD206^-^ (**Figure 1d**).

To test any potential link between BAT CX3CR1^+^ and CX3CR1^-^ myeloid cells, we administered TAM in *CX3CR1^CreERT^*^2^*^/GFP^; R26^LSL-tdT^* mice and followed tdTomato expression in BAT macrophages (**Figure 1e**). 2 days post-TAM administration, 66.6% of tdTomato^+^ BAT macrophages were CX3CR1^GFP+^ (**Figure 1e**). This percentage dropped to 33.8% 7 days after TAM (**Figure 1e**). We could thus demonstrate that monocytes are continuously recruited to BAT in which they first give rise to CX3CR1^+^ CD206^-^ CD226^-^ macrophages, that then differentiate to CX3CR1^-^ macrophages, corresponding to CD206^high^ and CD226^high^ cells.

The majority of blood monocytes arise from GMPs ^3^. To track GMP-dependent monocyte differentiation into BAT macrophages, we used *Ms4a3^CreERT^*^2^*; R26^LSL-tdT^* mice ^3^. Mice were gavaged with TAM and the frequency of tdTomato^+^ cells in each macrophage subset was defined in a time-course manner. As previously described ^3^, we observed that blood monocyte labelling efficiency reached 90-95% (data not shown). 3 days after the last TAM administration, around 70% of BAT monocytes were tdTomato^+^ (**Figure 1f**). This percentage rapidly decreased and reached 0% at day 17 due to the recruitment of unlabeled bone marrow-derived monocytes (**Figure 1f**). The labelling of DN BAT macrophages peaked at day 3, when 20% of cells were labelled, further supporting that this population depended on blood monocyte recruitment to sustain (**Figure 1f**). Of interest, the labelling of BAT CD226^high^ macrophages peaked at day 10 when labeling of DN macrophages was already decreasing (**Figure 1f**). This demonstrated that while CD226^high^ macrophages rely at least partially on GMP-derived monocytes for their maintenance, their differentiation trajectory occurs over a longer time frame. We detected very low contribution (less than 3%) of GMP-derived blood monocytes to BAT CD206^high^ macrophages (**Figure 1f**). However, it remains elusive whether early postnatal bone marrow-derived monocytes contribute to a long-lived pool of BAT-resident macrophages.

We next analyzed the presence of CD206^high^ and CD226^high^ at postnatal days (p)0, p7, p14, p21, and p28 in C57BL/6J mice. CD206^high^ macrophages were the only subset present at birth (p0), and CD226^high^ and DN macrophages only started to accumulate in BAT between p14 and p21 (**Figure S2a**). DN macrophages slowly accumulated over time, diluting CD206^high^ macrophages, while proportions of CD226^high^ macrophages remained constant (**Figure S2b**). To address the ontogeny and longevity of BAT macrophages at these pre- and postweaning stages, we utilized the “*double fate-mapper*” mouse model that enables the simultaneous labelling of macrophages derived from the yolk sac pre-macrophages (pMacs) and GMPs^18^. In this *double fate-mapper*, yolk sac-derived, tissue-resident macrophages express YFP via the *Tnfrsf11a^Cre^; Rosa26^LSL-YFP^* lineage tracer ^19^, while GMP-derived macrophages are labelled with tdTomato using the *Ms4a3^Flp^; Rosa26^FSF-tdT^* model (**Figure 1g**). The *double-fate mapper* also enables the distinction between short-lived (tdTomato^+^) and long-lived GMP-derived macrophages (YFP^+^ tdTomato^+^): as *Tnfrsf11a* is a macrophage core gene, its upregulation and concomitant expression of the Cre recombinase leads to YFP labelling of macrophages that take up tissue residency. YFP^+^ tdTomato^+^ CD206^high^ only started appearing at p14 at low frequencies (**Figure 1h**). At p21, we observed a transient influx of single tdTomato^+^ macrophages (∼8%), and a concurrent increase of YFP^+^ tdTomato^+^ CD206^high^ macrophages to 18%, indicating that weaning and/or food uptake leads to a recruitment of GMP-derived macrophages that become tissue-resident (**Figures 1g and S2c**). Albeit only very few CD226^high^ macrophages were detected at birth, all were single YFP^+^ (**Figures 1h and S2c**), suggesting that they develop from pMacs that were still circulating at birth and seeded the tissue in the first week of life. However, we observed a transient input of YFP^+^ tdTomato^+^ CD226^high^ macrophages (18%), similar to CD206^high^ macrophages. However, at p21, the GMP-derived, long-lived cells disappeared, and we detected an unlabeled CD226^high^ macrophage population (∼30%), indicating a GMP-independent influx of short-lived macrophages. Analysis of 8-week-old Ms4a3^Flp^ reporter mice showed that GMP contribution to BAT monocytes reached only ∼80%, compared to ∼95% in blood ^3^ (**Figures 1i-j**). This observation suggests that GMP-independent monocytes could be recruited to BAT. While 70% of DN macrophages were labeled, only 30% and 40% of CD226^high^ and CD206^high^ macrophages, respectively, were labeled (**Figure 1j**). In summary, these data demonstrate that all BAT-resident macrophages are embryonically-seeded, with the establishment of macrophage subset heterogeneity in the first weeks of life.

Lastly, we sought to define whether recruitment of monocytes is essential to the maintenance of BAT macrophage subsets. To answer this question, we treated mice with anti-CCR2 MC-21 antibody allowing to efficiently deplete classical monocytes ^20^. In BAT, we observed an almost complete monocyte depletion 24 hours after MC-21 treatment, in comparison to control animals (**Figure S2d**). Continuous MC-21 administration kept monocytes absent from BAT over time (**Figure 1k**). DN macrophage numbers tended to decrease at day 1 post MC-21 injection, while CD206^high^ and CD226^high^ macrophage abundance only started decreasing at day 2 post MC-21 administration (**Figure 1k**). Monocytes and all macrophage populations were decreased after 5 days of MC-21 treatment (**Figure 1k**). These data indicated that BAT macrophages strongly depend on monocyte flux to sustain constant numbers, albeit depletion of CD226^high^ and CD206^high^ macrophages was slower than DN cells (**Figure 1k**). Overall, our findings demonstrate the existence of robust monocyte contribution to BAT CD226^high^ macrophages through a minor GMP-dependent and a major GMP-independent cellular pathway.

### CD226^high^ macrophages preferentially arise from MDP-derived monocytes

A recent report suggested that a proportion of blood monocytes arise from a GMP-independent and MDP-dependent precursor ^15^. However, these observations relied on scRNA-seq data and adoptive transfer experiments, and genetic tools allowing to track MDPs and their progeny are lacking. We thus used RNA sequencing data from the Immgen consortium to compare GMPs to MDPs and identify subset-specific genes that could be targeted using existing genetic tools (**Figure S3a**). *Flt3* and *Ccr2* were present among the 30 most differentially expressed genes, however blood monocytes can be highly labeled using Flt3^Cre^ and Ccr2^Cre^ mice due to expression of these genes upstream and downstream of monocyte progenitors, making them incompatible with progenitor-restricted fate mapping. However, we identified *Clec9a* as a gene with higher expression in MDPs compared to GMPs (**Figure S3a**). Indeed, *Clec9a* appeared to be expressed in MDPs and CDPs but not in upstream progenitors nor GMPs and monocytes (**Figure 2a**). We thus crossed Clec9a^iCre^ mice ^21^ to R26^TdT^ mice and analyzed bone marrow progenitor labelling efficiency (**Figure S3b**). Our data revealed that 40% of CDPs and 10% of MDPs were tdTomato^+^ (**Figure 2b**). cMoPs were not labelled and remained tdTomato^-^ (**Figure 2b**). We also detected a population of bone marrow monocytes (around 5%) that expressed tdTomato (**Figure 2b**). These data indicated that Clec9a labelling in monocytes could reflect a MDP-dependent origin.

**Figure 2.**
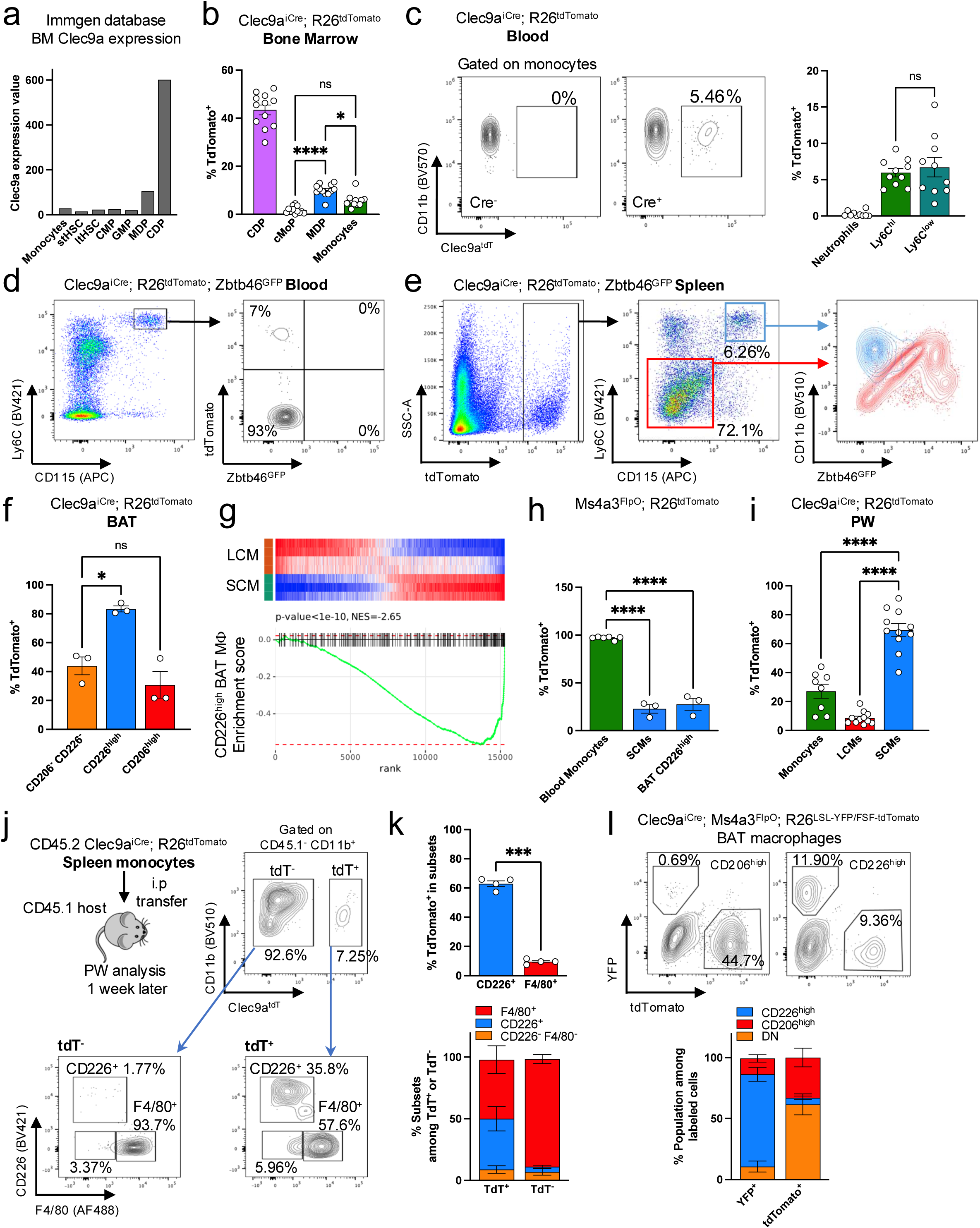
CD226^high^ macrophages develop from MDP-derived monocytes. (**a**) Expression of *Clec9a* mRNA by bone marrow (BM) short term hematopoietic stem cells (stHSC), long term hematopoietic stem cells (ltHSC), common myeloid progenitors (CMP), granulocyte monocyte progenitors (GMP), monocyte dendritic cell progenitors (MDP), common dendritic cell progenitors (CDP) and monocytes. Data from Immgen.org. (**b**) Quantification of tdTomato^+^ cells am ong bone marrow progenitors and monocytes in Clec9a^iCre^; R26^tdTomato^ mice. (**c**) Quantification of tdTomato^+^ cells am ong blood neutrophils and monocytes in Clec9a^iCre^; R26^tdTomato^ mice. (**d-e**) Flow cytometry analysis of blood (d) and spleen (e) Ly6C^high^ monocytes from Clec9a^iCre^; R26^tdTomato^; Zbtb46^GFP^ mice. (**f**) Quantification of tdTomato^+^ cells am ong BAT macrophages in Clec9a^iCre^; R26^tdTomato^ mice. (**g**) GSEA analysis, comparing the signature of BAT CD226^high^ macrophages to peritoneal LCMs and SCMs. (**h**) Quantification of tdTomato^+^ cells am ong blood monocytes, peritoneal SCMs and BAT CD226^high^ macrophages in Ms4a3^FlpO^; R26^tdTomato^ mice. (**i**) Quantification of tdTomato^+^ cells am ong peritoneal monocytes and macrophages in Clec9a^iCre^; R26^tdT^ mice. (**j**) Experimental design and representative flow cytometry plots showing phenotype of tdTomato^+^ and tdTomato^-^ monocytes from Clec9a^iCre^; R26^tdT^ mice that were adoptively transferred intraperitoneally. (**k**) Quantification of tdTomato^+^ cells am ong peritoneal wash F4/80^+^ and CD226^high^ cells following adoptive transfer as in (j). (**l**) Representative flow cytometry plots showing YFP and tdTomato expression am ong BAT macrophages (top) and proportions of macrophage subsets am ong YFP^+^ or TdTomato^+^ cells (bottom) from Clec9a^iCre^; Ms4a3^FlpO^; R26^LSL-YFP/FSF-tdTomato^ mice. Data are presented as mean values +/- SEM. See also Figure S3.

We next analyzed the presence of tdTomato^+^ cells in peripheral blood immune cell populations in Clec9a^iCre^ R26^tdT^ mice (**Figure S3c**), demonstrating that approximately 5% of blood monocytes were labelled (**Figure 2d**), mirroring blood monocyte labeling using Ms4a3^Flp^ mice. Equal tdTomato labeling was observed in Ly6C^high^ and Ly6C^low^ blood monocytes (**Figure 2d**). However, no additional blood immune cells were labelled. Indeed, neutrophils, B cells, T cells (both CD4^+^ and CD8^+^ subsets) and Natural Killer (NK) cells remained tdTomato^-^ (**Figure S3d**). To verify whether tdTomato^+^ blood CD11b^+^ CD115^+^ Ly6C^high^ cells were indeed monocytes rather than pre-DCs, we crossed Clec9a^iCre^ R26^TdT^ mice with Zbtb46^GFP^ reporter mice. We confirmed that tdTomato expression was present in bone marrow CD11b^-^ CD34^+^ CD16/32^-^ Flt3^+^ cKit^+^ Zbtb46^GFP-^ progenitors, containing MDPs (**Figure S3e**). CD11b^-^ CD34^+^ CD16/32^-^ Flt3^+^ cKit^-^ Zbtb46^GFP+^ progenitors, containing CDPs, also highly expressed tdTomato (**Figure S3e**). In blood, both tdTomato^+^ and tdTomato^-^ Ly6C^high^ monocytes lacked GFP signal (**Figure 2d**). We then analyzed the spleen since this tissue contains DCs that could act as internal positive control for GFP expression (**Figure 2e**). Ly6C^high^ monocytes constituted around 6% of tdTomato^+^ cells in this organ, and again these cells lacked expression of Zbtb46^GFP^ (**Figure 2e**). We next sought to analyze Clec9a^iCre^ labeling among tissue macrophages and observed that the majority (70-80%) of BAT CD226^high^ macrophages were labelled with tdTomato (**Figure 2f**). These data mirrored the labelling percentage documented in Ms4a3^Flp^; R26^tdTomato^ mice, suggesting that MDP-derived monocytes rather than GMP-derived monocytes could be the main progenitors of CD226^high^ macrophages.

We next asked whether a similar labeling pattern could be observed in macrophages from other tissues. Small peritoneal cavity macrophages (SCMs) were demonstrated to express CD226 and to heavily rely on monocytes for their maintenance ^22^. These cells reside in the peritoneal cavity where they share the local microenvironment with large peritoneal cavity macrophages (LCMs), yet LCMs and SCMs display specific transcriptional signatures (Immgen.org) (**Figure 2g**). We compared the transcriptomic profile of BAT CD226^high^ macrophages to that of peritoneal LCMs and SCMs and found that they strongly resembled SCMs (**Figure 2g**), indicating that the CD226^high^ macrophage subset could be conserved across several tissues. Peritoneal wash (PW) SCMs and BAT CD226^high^ macrophages showed a similar labeling pattern in Ms4a3^Flp^; R26^tdTomato^ mice with only around 25% of these cells being labeled (**Figure 2h**). Curiously, around 20% of monocytes were labeled in PW from Clec9a^iCre^; R26^tdTomato^ mice (**Figure 2i**) compared to 5% in blood. While only a small fraction of LCMs was tdTomato^+^, around 75% of SCMs expressed the tdTomato reporter (**Figure 2i**). We confirmed that this was not due to *Clec9a* mRNA expression in SCMs (**Figure S3f**) nor BAT CD226^high^ macrophages (**Figure S3g**). Taken together, these data suggested that MDP-derived monocytes give preferentially rise to peritoneal cavity and BAT CD226^high^ macrophages.

Based on the observation that MDP-derived monocytes had a higher contribution to CD226^high^ macrophages than GMP-derived monocytes across tissues, we hypothesized that monocyte differentiation trajectories were pre-conditioned by their origin. To test this hypothesis, we cell-sorted total splenic monocytes from Clec9a^iCre^ R26^TdT^ mice (CD45.2^+^ cells) and transferred them intraperitoneally into CD45.1 host mice without prior treatment (**Figure 2j**). 7 days after injection, the presence and phenotype of donor CD45.2 cells were analyzed by flow cytometry. We observed that tdTomato^-^ donor cells preferentially gave rise to LCMs (**Figures 2j and 2k**). In striking contrast, tdTomato^+^ cells generated both CD226^high^ SCMs and LCMs (**Figures 2j and 2k**). Even though only ∼7% of donor cells were tdTomato^+^ (**Figure 2j**), 60% of donor CD226^high^ SCMs were tdTomato^+^ (**Figure 2k**). These results demonstrate that monocyte origin influences their ability to generate specific macrophage subsets in tissues.

Finally, we sought to confirm that Clec9a^iCre^ and Ms4a3^Flp^ mice allowed to label two different lineages. For this purpose, we crossed Ms4a3^Flp^; R26^FSF-tdTomato^ mice Clec9a^iCre^ R26^LSL-YFP^ mice to generate a double reporter system in which MDP-derived macrophages would be YFP-labeled and GMP-derived macrophages would be tdTomato-labeled (**Figure 2l**). We observed non-overlapping expression of YFP and tdTomato among BAT CD226^high^ macrophages (**Figure 2l**). Even though overall YFP labeling was lesser than what we measured using Ai14 reporter mice (**Figure S3h**) as expected, most YFP^+^ cells were CD226^high^ and most tdTomato^+^ cells were CD206^high^ or CD206^-^CD226^-^ macrophages in BAT (**Figure 2l**). Both YFP and tdTomato labeling were very low or absent in resident macrophage populations such as alveolar macrophages, Kupffer cells and microglia (**Figure S3i**). Overall, our data validate Clec9a^iCre^ mice as suitable tools for fate mapping MDP-derived monocytes and macrophages and demonstrate that CD226^high^ macrophages are a population with mixed monocytic origin dominated by the MDP lineage.

### CD226^high^ macrophages are conserved across tissues and enriched in thermogenic adipose depots

We observed that a myeloid cluster also expressed CD226 in our previously published single cell RNA-sequencing dataset of adrenal gland (AG) immune cells ^23^. A recent direct comparison between murine and human peritoneal cavity immune cells revealed that CD226^high^ SCMs are conserved in humans and show similarities with human type 2 dendritic cells ^24^. Based on these observations, we aimed to decipher (i) whether CD226^high^ macrophages were present in other tissues beyond BAT and peritoneal cavity and (ii) whether these cells might share part of their signature with dendritic cells. To answer these questions, we integrated mononuclear phagocytes from previously published scRNA-seq datasets from BAT ^5^, AG ^23^ and peritoneal cavity ^24^, as well as splenic Flt3^+^ cells containing plasmacytoid and conventional (type 1, 2 and 3) dendritic cells ^25^ (**Figure 3a**). CD226^+^ macrophages were identified in BAT, AG and peritoneal cavity and they clustered together (**Figures 3a-b**). Importantly, CD226^high^ cells clustered away from recently-described type 3 dendritic cells (cDC3) that express markers common with the macrophage lineage including CD16/32 and CD115 ^25^ (**Figure 3b**). Indeed, CD226^high^ macrophages lacked expression of the core signature of cDC3 (**Figure 3c**). We next explored a publicly available dataset generated by the Tabula Muris consortium ^26^ to test whether CD226^+^ macrophages could be found in additional tissues. We generated a gene set named “CD226^high^ signature”, comprising the 50 genes with highest differential expression between BAT CD226^high^ macrophages and other BAT immune cells. This signature was absent in most of the dataset, except for one group of cells (**Figure S4a, left**). Indeed, cells highly expressing this gene set clustered together (**Figure S4a, right**). These cells were predicted to be myeloid cells and in particular macrophages (**Figure S4b**). They were enriched in adipose tissue and limb muscle, even though they were also detected at a lower degree in heart, trachea and mammary glands (**Figure S4b**). In agreement with our data presented in Figure S2a, the abundance of CD226^high^ macrophages did not depend on biological sex nor age (**Figure S4b**).

**Figure 3.**
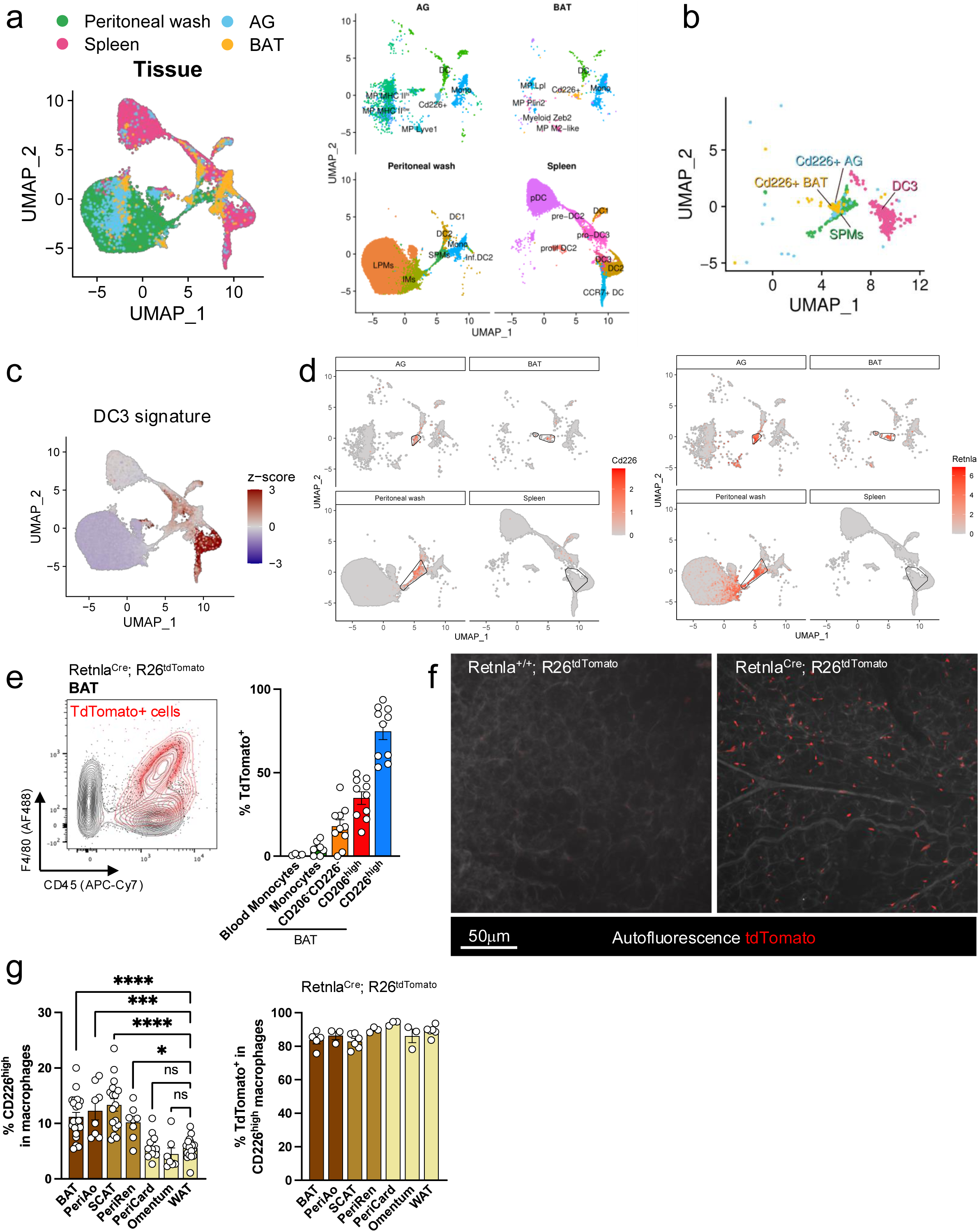
CD226^high^ macrophages are conserved across tissues and enriched in thermogenic adipose tissue. (**a**) Single cell RNA-sequencing analysis of integrated datasets of immune cells from BAT, adrenal glands (AG) and peritoneal wash, and Flt3^+^ cells from spleen. (**b**) UMAP plot representing clustering of CD226^high^ macrophages from BAT, AGs and peritoneal wash, and DC3 from spleen. (**c**) UMAP plot representing expression of the DC3 signature by all cells contained in the integrated dataset from (a). (**d**) Expression of *Cd226* and *Retnla* by BAT, AG, peritoneal wash and spleen immune cells. (**e**) Flow cytometry plot showing distribution of tdTomato^+^ cells from Retnla^cre^; R26^tdTomato^ mice in BAT (left) and quantification of tdTomato expression by blood and BAT monocytes and BAT macrophages (right). (**f**) Confocal microscopy analysis of BAT from Retnla^+/+^; R26^tdTomato^ and Retnla^Cre/+^; R26^tdTomato^ mice. (**g**) Quantification of F4/80^+^ CD64^low^ CD226^high^ macrophages (left) and their expression of tdTomato in Retnla^cre^; R26^tdTomato^ mice (right) in interscapular BAT, peri-aortic brown adipose tissue (PeriAo), subcutaneous adipose tissue (SCAT), peri-renal adipose tissue (PeriRen), peri-cardiac adipose tissue (PeriCard), omentum and gonadal white adipose tissue (WAT). Data are presented as mean values +/- SEM. See also Figure S4.

Next, we generated a genetic model to target and track CD226^+^ macrophages across tissues. Resistin like alpha (*Retnla*) appeared to be a hallmark gene for both BAT and peritoneal CD226^+^ macrophages ^5,22^. We could confirm that *Retnla* was highly expressed in CD226^high^ macrophages in our integrated dataset (**Figure 3d**). Thus, we crossed Retnla^Cre^ mice ^27^ with *R26^LSL-tdTomato^* reporter animals. In these mice, tdTomato expression was exclusively detected in CD45^+^ cells in BAT (**Figure 3e**). Among CD45^+^ cells, tdTomato expression was highly enriched in myeloid F4/80^+^ cells (**Figure 3e**). Among BAT macrophages, the highest tdTomato expression was detected in CD226^high^ macrophages, although lower labelling was also detected in BAT CD206^high^ and DN macrophages (**Figure 3e**). Blood monocytes were not labeled with tdTomato (**Figure 3e**). We could confirm that only cells resembling macrophages expressed tdTomato in BAT by confocal microscopy, and that adipocytes were not labelled (**Figure 3f**). In the peritoneal cavity, we found that tdTomato was expressed by SCMs but not by dendritic cells (**Figure S4c**). We next investigated the presence of CD226^high^ macrophages in different adipose tissue depots and compared brown (BAT, periaortic AT (PeriAo)), beige (subcutaneous (SCAT) and perirenal (PeriRen) AT) and white (pericardiac AT (PeriCard), omentum and gonadal WAT) fat, all associated with different thermogenic capacity, inferred on adipocyte phenotype (**Figure S4d**). While CD226^high^ macrophages were detected in all these tissues, their relative frequency was increased within tissues containing multilocular adipocytes (**Figure 3g**). Furthermore, these cells were efficiently labelled in Retnla^Cre^; R26^tdTomato^ mice (**Figure 3g**). Together, our data indicate that CD226^high^ Retnla^+^ myeloid cells are a unique macrophage subset that is different from conventional dendritic cells, conserved across tissues and enriched in thermogenic adipose depots.

### CD226^high^ macrophages are regulated by CSF1R and GM-CSF

We next aimed to identify growth factors regulating CD226^high^ macrophages. We could detect differential expression of *Csf1r* and *Csf2ra* mRNA between CD206^high^ and CD226^high^ macrophages in BAT scRNA-sequencing data ^5^ (**Figure 4a**), suggesting that these cells might be regulated by different growth factors. Similar observations were made when analyzing peritoneal cavity SCMs and LCMs transcriptomic signatures (Immgen.org) (**Figure S5a**). First, we treated mice with anti-CSF1R blocking Ab (clone AFS98). In BAT, AFS98 administration did not induce a complete macrophage depletion (**Figure 4b**). On the level of macrophage subsets, AFS98 administration decreased the proportions and numbers of BAT CD206^high^ macrophages (**Figures 4c-d**). CD226^high^ cells were also negatively impacted, although at a lower magnitude compared to the CD206^high^ subset (**Figure 4d**) resulting in increased proportions of CD226^high^ cells (**Figure 4c**). Similarly, in the peritoneal cavity AFS98 treatment led to a 70% reduction in LCM numbers while SCM numbers were reduced by 47% indicating a lesser dependence on CSF1R compared to LCMs (**Figure S5b**). Thus, in BAT CD206^high^ macrophages appeared to be the population with the highest dependence on CSF1R signaling.

**Figure 4.**
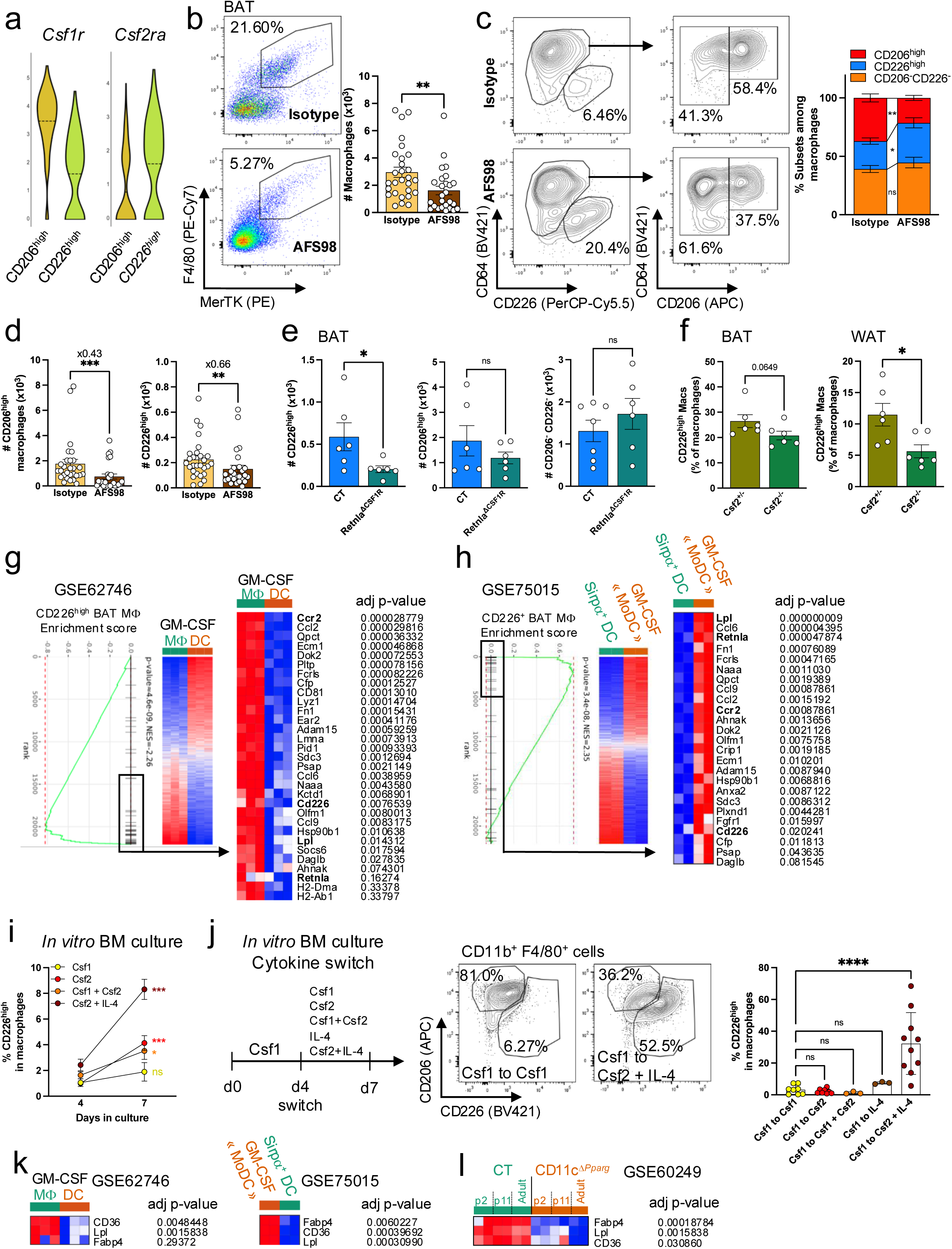
CSF1R and GM-CSF regulate CD226^high^ macrophages. (**a**) Violin plots representing *Cs1r* and *Csf2ra* expression by BAT CD226^high^ and CD206^high^ macrophages. (**b-d**) Representative flow cytometry plots and quantifications of total BAT macrophages (b) and BAT macrophage subset numbers (c) and proportions (d) following CD115 blockade using AFS98. (**e**) Quantification of BAT macrophage susbets in control (CT) and Retnla^ΔCSF1R^ mice. (**f**) Quantification of BAT macrophage subsets in Csf2^+/-^ and Csf2^-/-^mice. (**g, h**) GSEA analysis and differential gene expression analysis, comparing the signature of BAT CD226^high^ macrophages to macrophages and DCs obtained by culturing bone marrow with GM-CSF (g) or conventional Sirpα^+^ DCs and “MoDCs” obtained with GM-CSF culture (h). (**i**) Flow cytometry analysis of total bone marrow cells cultured with M-CSF (50ng/m L), GM-CSF (50ng/m L) with or without IL-4 (10ng/m L), or M-CSF + GM-CSF (50ng/m L) for 4 or 7 days. (**j**) Flow cytometry analysis of total bone marrow cells cultured with M-CSF for 4 days, and then with M-CSF, GM-CSF with or without IL-4, or M-CSF + GM-CSF for another 3 days.(**k, l**) Expression of *Cd36*, *Fabp4* and *Lpl* measured in GM-CSF derived macrophages and dendritic cells (k) or alveolar macrophages from new born (p2, p11) or adult (7-8 weeks) control (CT) and CD11^cre^; Pparg^fl/fl^ mice (l). Data are presented as mean values +/- SEM. See also Figure S5.

To test whether the reduction of CD226^high^ macrophages induced by AFS98 treatment was cell autonomous or was tied to the disappearance of CD206^high^ macrophages, we generated Retnla^Cre^ x Csf1r^fl/fl^ (Retnla^ΔCSF1R^) mice. In Retnla^ΔCSF1R^ mice, we could observe a two-fold reduction of CD226^high^ macrophage numbers in BAT, SCAT, WAT and PW while other populations were unaffected (**Figures 4e and S5c-d**). Yet, this reduction did not align with Retnla^Cre^ efficiency in this population that reached ∼85-90% (**Figure 2g**), suggesting that cells remaining in Retnla^ΔCSF1R^ mice were not CSF1R-dependent.

Based on our transcriptomic observations (**Figure 4a**), we next sought to test whether GM-CSF could be regulating CD226^high^ macrophages. CD226^high^ macrophages were recently observed in exocrine glands, in which they were called adenophages and in which they rely on GM-CSF produced by ILC2 ^28^. To test whether such a mechanism could be conserved in adipose tissue, we analyzed GM-CSF fate mapping and reporter FROGxAi14 mice that label present (GFP^+^ tdTomato^+^) and past (GFP^-^ tdTomato^+^) GM-CSF-producing cells ^29^. In WAT, ∼25% of CD45^+^ cells were tdTomato^+^ and ∼10% were tdTomato^+^ GFP^+^ (**Figure S5e**). Among GFP^+^ cells, ∼75% were ILC2 (**Figure S5h**). To test whether GM-CSF regulated CD226^high^ macrophages, we next analyzed double knock-in/knock-out GM-CSF deficient Csf2^LacZ^/^LacZ^ (Csf2^-/-^) mice ^28^. Similarly to Retnla^ΔCSF1R^ mice, CD226^high^ macrophages were partially impacted in Csf2^-/-^ mice (**Figure 4f**). Curiously, in peritoneal wash this phenotype was only observed in male Csf2^-/-^ mice (**Figure S5f**).

Culturing bone marrow cells *in vitro* with GM-CSF yields both macrophages (GM-Macs) and dendritic cells (GM-DCs) ^30^. To test whether the signature of BAT CD226^high^ macrophages overlapped with either GM-CSF-generated cell types, we performed GSEA analysis (**Figures 4g and S5g**). We found that hallmark genes of BAT CD226^high^ macrophages were mainly expressed in GM-Macs, including *Cd226* and *Fn1* (**Figure 4g**). Similarly, Ly6C^high^ monocytes have been shown to generate “Mo-DCs” upon culture with GM-CSF ^31^. Again, we found that the BAT CD226^high^ macrophage signature aligned with Mo-DCs rather than Sirpα^+^ conventional dendritic cells (**Figure 4h**). Although BAT CD226^high^ macrophage signature tended to be regulated when GM-CSF treatment was coupled with IL-4, no statistically significant differences were observed (**Figure S5h**). We repeated such *in vitro* experiments to validate the appearance of CD226^high^ macrophages by flow cytometry. While culturing bone marrow cells with only Csf1 for one week did not favor the generation of these cells, presence of Csf2 triggered development of CD226^high^ macrophages and this phenotype was exacerbated by addition of IL-4 (**Figure 4i**). We then asked whether exposure to these cytokines could skew differentiating macrophages towards the CD226^high^ macrophage phenotype. To test this hypothesis, bone marrow cells were cultured with Csf1 for 4 days, then washed with PBS and cultured for another 3 days in media containing Csf1, Csf2, Csf1 + Csf2, IL-4 or Csf2 + IL-4 (**Figure 4j**). Proportions of CD226^high^ macrophages augmented > 10-fold in presence of Csf2 and IL-4, while other conditions did not trigger such an effect (**Figure 4j**).

BAT macrophages express a clear lipid-associated signature, including *Cd36*, *Fabp4* and *Lpl* ^5^, with the latter being a core signature gene of CD226^high^ macrophages (**Figures 4g-h**) and of closely-related immature CD206^-^CD226^-^*Lpl*^high^ macrophages (**Figure 1a**). GM-CSF-derived macrophages highly expressed these three genes in comparison with their DC counterparts (**Figure 4k**). Since *Cd36*, *Fabp4* and *Lpl* are PPARγ-induced genes, we analyzed their expression in GM-CSF-dependent alveolar macrophages ^32^ and could confirm significant downregulation in *Pparg*-deficient alveolar macrophages (**Figure 4l**), suggesting that GM-CSF could be regulating CD226^high^ macrophages through PPARγ.

Together, our results indicate that IL-4 and several growth factor-dependent pathways, notably GM-CSF and CSF1R-mediated signaling, regulate CD226^high^ macrophages. GM-CSF could regulate CD226^high^ macrophage expression of lipid-associated genes like *Lpl*, *CD36* and *Fabp4*. The partial phenotypes observed in Retnla^ΔCSF1R^ and Csf2^-/-^ mice could reflect the complementarity of Csf1 and Csf2 observed in vitro (**Figure 4j**), or perhaps the ontogenic heterogeneity of CD226^high^ macrophages (**Figure 2**). Whether M-CSF and GM-CSF differently affect monocyte generation and differentiation depending on their origin remains to be determined.

### CD226^high^ adipose tissue macrophages regulate lipid metabolism

We next sought to investigate the role of BAT macrophage subsets on tissue lipid content. To target highly CSF1R-dependent CD206^high^ macrophages, we repeated our AFS98-mediated depletion approach. Mouse body weight and BAT weight tended to decrease in AFS98-injected mice in comparison to isotype controls (**Figure S6a**). AFS98 triggered a well-defined morphological change in BAT with diminished presence of adipocytes with large lipid droplets in comparison to control treated animals (**Figure 5a**). Enzymatic triglyceride (TG) dosage confirmed that BAT TG content was decreased in AFS98 injected mice (**Figure 5b**). Of note, serum TG levels remained similar between both groups of animals (**Figure S6b**). Free fatty acids concentration and diversity were not modified by AFS98 administration (**Figure S6c**). These results indicate that CD206^high^ macrophages could support triglyceride storage in BAT, a finding in agreement with recently published results in WAT ^9^.

**Figure 5.**
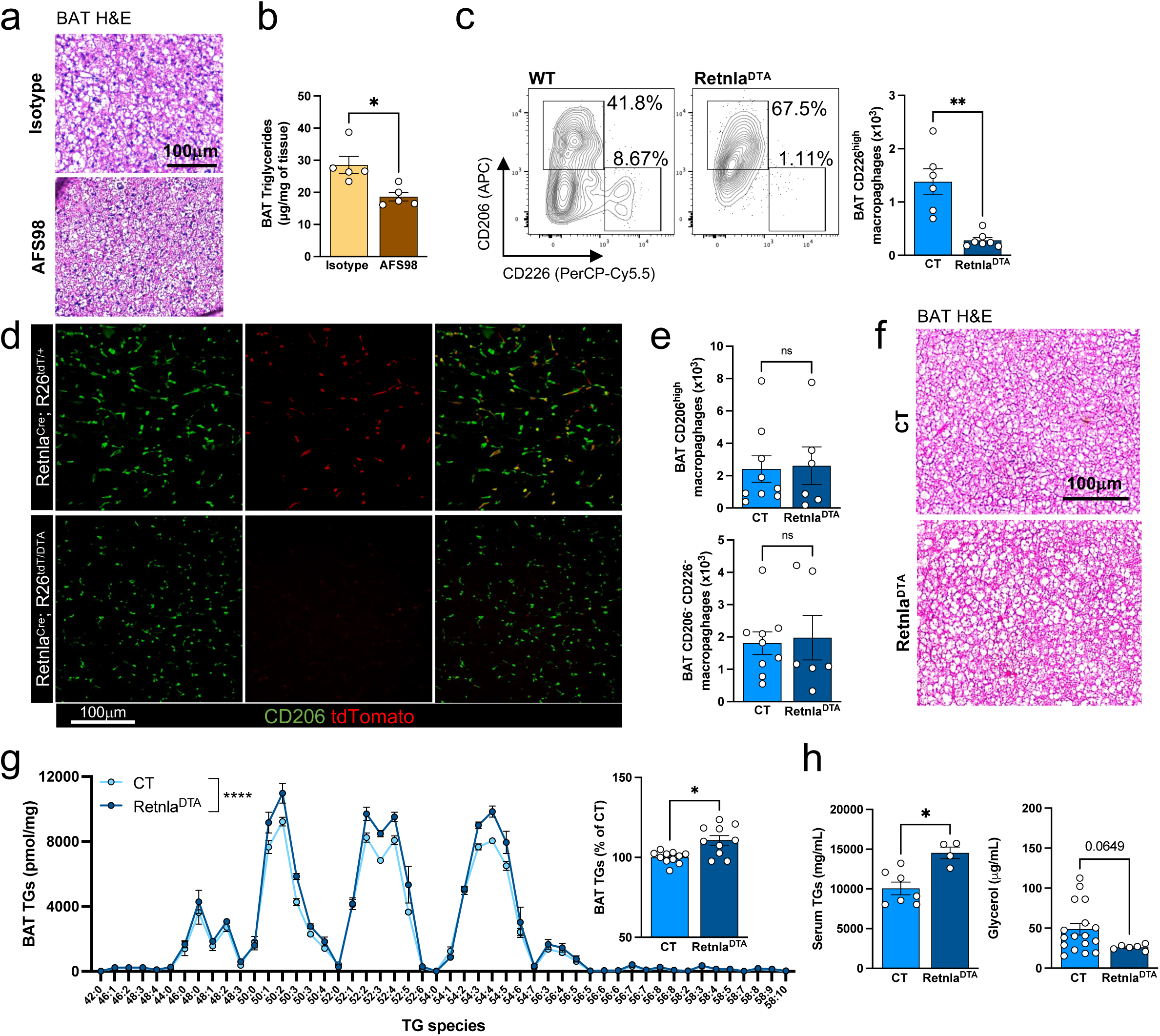
CD226^high^ macrophages regulate triglyceride metabolism. (**a-b**) Hematoxylin and eosin staining of BAT (a) and quantification of BAT triglyceride content (b) following AFS98 treatment. (**c**) Representative flow cytometry plots (left) and quantification (right) of BAT CD226^high^ macrophages in control (CT) and Retnla^Cre^; R26^DTA^ mice. (**d**) Confocal microscopy analysis of BAT from Retnla^Cre^; R26^tdT/DTA^ and control Retnla^Cre^; R26^tdT/+^ mice. (**e**) Representative flow cytometry plots (left) and quantification (right) of BAT CD206^high^ and CD206^-^ CD226^-^ macrophages in control and Retnla^Cre^; R26^DTA^ mice. (**f, g**) Hematoxylin and eosin staining of BAT (e) and triglyceride content analyzed using lipidomics (f) in Retnla^Cre^; R26^DTA^ mice. (**g**) Quantification serum triglyceride (TG) and glycerol levels in Retnla^Cre^; R26^DTA^ mice. Data are presented as mean values +/- SEM. See also Figure S6.

Since AFS98 treatment could also partly affect CD226^high^ macrophages, we next investigated the role of these cells using Retnla^Cre^ mice. We generated Retnla^Cre^ R26^DTA^ mice (Retnla^DTA^) to deplete CD226^high^ macrophages across tissues. Blood immune cells were not affected in these mice, indicating the absence of systemic inflammation (**Figure S6d**). However, we could confirm an almost complete absence of BAT Retnla^+^ CD226^high^ macrophages by flow cytometry and confocal microscopy (**Figures 5c-d**). Importantly, CD206^high^ and DN BAT macrophage numbers remained similar between control and Retnla^DTA^ animals (**Figure 5e**). We also noticed the disappearance of CD226^high^ macrophages in all adipose tissue depots that we analyzed (**Figure S6e)**. We also noticed a complete disappearance of SCMs in the peritoneal cavity while LCMs were not affected (**Figure S6f**). Other immune cell types were also unaffected in BAT (**Figure S6g**). Retnla^DTA^ mice are thus appropriate tools to address the function of CD226^high^ macrophages.

Depletion of CD226^high^ macrophages triggered a specific BAT phenotype characterized by increased adipocyte droplet size (**Figure 5f**). This was associated with an increased total BAT TG content in Retnla^DTA^ mice when compared to littermate controls, with a particular enrichment of long-chain TGs (**Figure 5g**). We did not detect changes in serum non esterified fatty acids (NEFA) and glucose levels in Retnla^DTA^ mice (**Figure S6h**). However, their serum TG levels were higher than controls while their glycerol levels trended downwards (**Figure 5h**), indicating that CD226^high^ macrophages were regulating triglyceride metabolism.

## Discussion

In the present study, using genetic fate mapping, we demonstrated that a population of blood monocytes is generated independently of GMP-derived cells. We found that these cells are efficiently labelled in Clec9a^iCre^ R26^TdT^ mice suggesting that those monocytes arise mainly from MDPs. Of note, these monocytes preferentially generated CD226^high^ macrophages in adipose tissue and peritoneal cavity, demonstrating the monocyte origin influences their fate. Whether other macrophage subsets also arise mainly from MDPs remains to be established. While the phenotypic distinction between monocytes, macrophages and DCs was challenging for a long time, key findings reported in 2012 by the Immgen consortium solved this problem ^33,34^. Macrophages were defined as CD64^+^MerTK^+^ cells, markers that were not expressed on conventional DCs ^33^. We found that CD226^high^ macrophages across tissues displayed a low CD64 expression, challenging the current classification of cells in the macrophage or DC lineage based on these criteria. While BAT CD226^+^ macrophages expressed MerTK and F4/80, supporting their macrophage identity, they also expressed CD11c and MHC-II, two markers traditionally detected and associated with DCs. A key feature of DCs, that is not shared with tissue resident macrophages, is their ability to migrate through the lymphatic vessels to local draining lymph nodes ^35^. DC migration via the lymphatic endothelial system depends on a single chemokine receptor, CCR7. Indeed, CCR7 expression allows to discriminate between DCs and macrophages. CD226^high^ macrophages lack CCR7 expression (data not shown), thus suggesting that these cells cannot migrate through the lymphatic vessels. Furthermore, our data revealed that GMP- and MDP-derived monocytes contribute both to the generation of CD226^high^ macrophages. Therefore, one could expect that depending on their origin, GMP- and MDP-derived CD226^high^ macrophages could display selective functions. Their high MHC-II expression suggests a role in antigen presentation and initiation of adaptive immunity. The ability of GMP- and MDP-derived CD226^high^ macrophages to trigger a T cell response remains to be established.

Previous work on SCMs demonstrated that these cells rely on the transcription factor IRF4 ^22^. This study indicated that, independently of their origin, SCMs rely on a unique transcription factor for their survival. However, while only 5% of blood monocytes originate from MDPs, 70% of SCMs arose from these monocytes. Monocytes rely on the chemokine receptor CCR2 for their bone marrow egress and tissue entry ^36^. While SCM numbers are reduced in CCR2-deficient mice ^22^, they are not fully abolished. One could wonder whether MDP-derived monocytes rely solely on CCR2 for bone marrow egress and their subsequent tissue recruitment. Mechanisms governing MDP-derived monocyte bone marrow retention and egress thus require further investigations that are beyond the focus of the present study. We observed a conserved ratio of Ly6C^high^ : Ly6C^low^ monocytes independently of their origin, suggesting that monocyte conversion occurs at a similar rate in GMP- and MDP-derived monocytes. Yet, our data indicates that GMP- and MDP-derived monocyte fate in tissues depends on their lineage. MDP-derived monocytes are preferentially recruited to specific tissues, i.e. adipose tissues and limb muscle where CD226^+^ macrophages are enriched. A recent study suggests that monocytes locally proliferate upon their tissue recruitment before giving rise to tissue resident macrophage subsets ^37^. The proliferative rate of GMP- and MDP-derived monocytes could be significantly different, and this might explain, at least partially, their ability to differentially contribute to specific tissue resident macrophage subsets. The lifespan of GMP- and MDP-derived CD226^high^ macrophages could be an additional factor involved in the aforementioned process.

CD226^high^ macrophages were enriched in adipose tissue and in particular thermogenic depots. Depletion of CD226^high^ macrophages was associated with an increased local and systemic TG content, demonstrating a central role of these cells in lipid metabolism. While CD226^high^ macrophages express several enzymes involved in lipid metabolism ^5^, how precisely these cells control local and systemic TG content requires further investigations. Previous reports demonstrated that adipose tissue macrophages control lipid storage in a PDGFcc-dependent manner ^9^. Of note, mice treated with anti-CSF1R blocking antibody and CCR2^-/-^mice displayed a different behavior when challenged with high fat diet ^9^. While both CD206^high^ and CD226^high^ macrophages show CCR2-dependent monocyte contribution (Figure 1) ^5^, monocytes mainly contribute to the DN and CD226^high^ subsets. Our data demonstrated that adipose tissue CD206^+^ macrophages showed a higher CSF1R dependency in comparison to DN and CD226^high^ macrophages. Using our scRNA-seq data, we could determine that *Csf1r* expression is higher in CD206^high^ macrophages compared to the CD226^high^ population (data not shown), which could explain the differential impact of AFS98 administration between these two subsets. Whether GMP- and MDP-derived CD226^+^ macrophages rely to a similar extent on CSF1R signaling for their proliferation and renewal needs to be defined. Inversely, CD226^high^ macrophages expressed high levels of *Csf2ra* mRNA compared to CD206^high^ macrophages (data not shown). This observation suggests that GM-CSF signaling plays a role in the maintenance of CD226^high^ macrophages.

In the present study, we identify CD226^high^ macrophages as a new myeloid subset bearing features of both macrophages (F4/80^+^ MerTK^+^) and dendritic cells (CD64^low^ CD11c^+^ MHC-II^high^), that is conserved across tissues. Similar findings were reported in a recent study comparing murine and human peritoneal cells ^24^. Indeed, the human counterpart to murine CD226^high^ SCMs appeared to be CD14^+^ cells that also express the DC marker CD1c ^24^. Importantly, this observation indicates that CD226^high^ cells described herein are conserved in humans. We detected a similar population in human white adipose tissue (Human Single Cell Atlas from the Broad Institute, data not shown), indicating that these cells are also conserved across several tissues in humans. This work paves the way to define the relative role of GMP-and MDP-derived monocytes during infections, acute and chronic (obesity, atherosclerosis…) inflammatory diseases and cancer development.

## Acknowledgments

We thank the C3M Animal facility, and the C3M Flow Cytometry Core Facility financed by Conseil Général CG06 and Conseil Régional PACA. We thank Dr. Florent Ginhoux for sharing genetic mouse models. We thank Sandrine Quemener (Lille Pasteur Institute) for technical assistance. SI is funded by Institut National de la Sante et de la Recherche Medicale (INSERM) and Agence Nationale de la Recherche (ANR-22-CE14-0027-02; ANR-21-CE15-0020-02; ANR-23-CE14-0054-02; ANR-23-CE15-0032-01). JWW was supported by NIH R01AI165553. MC was supported by NIH T32 HL166142. AG was supported by the French government through the France 2030 investment plan managed by the National Research Agency (ANR), as part of the Initiative of Excellence Université Côte d’Azur under reference number ANR- 15-IDEX-01. EM was funded by the Deutsche Forschungsgemeinschaft (DFG, German Research Foundation) under Germany’s Excellence Strategy-EXC2151-390873048, SFB1454 (Project number 432325352), and FOR5547 – Project-ID 503306912.

## Author contribution

AG and SI designed the study. AG and SI wrote the manuscript and prepared the figures with input from all co-authors. AG, ZC, FW, ST, SG, MT, AC, LL, KA, LL, MG, TP, SF, MC, BD, EG, MF, EB, ZM, FNZ, JGN, JWW, AB and SI performed experiments. AG, ZC, FW, ST, MT, KA, SG, TP, JWW, DM, MNA, AB and SI analyzed data. EM, DV, MM, JH, DD and BB provided tools and expertise. SI obtained funding for the project.

## Methods

### Experimental models

#### Mouse models

Wild-type C57BL/6J (Jax #000664), CX3CR1^gfp^ (B6.Cg-Ptprca Cx3cr1^tm1Litt/LittJ^, Jax #008451), Ms4a3^cre^ (C57BL/6J-Ms4a3^em2(cre)Fgnx^/J, Jax #036382), R26^LSL-tdTomato^ (B6.Cg-Gt(ROSA)26Sor^tm9(CAG–tdTomato)Hze^/J, Jax#007909), Clec9a^icre^ (B6J.B6N(Cg)-Clec9a^tm2.1(icre)Crs^/J, Jax#025523), CD45.1 (B6.SJL-Ptprc^a^ Pepc^b^/BoyJ, Jax# 002014) and R26^LSL-DTA^ (B6.129P2-Gt(ROSA)26Sor^tm1(DTA)Lky^/J, Jax# 009669) mice used in this study were maintained under C57BL/6J background and originally purchased from Jax Labs. CCR2^cre/ERT2^ (C57BL/6NTac-Ccr2^tm2982(T2A-Cre7ESR1-T2A-mKate2^) ^38^ and Csf2^-/- 28^ mice were provided by Dr. Burkhard Becher.

Ms4a3^cre/ERT2^ mice^3^ were provided by Dr. Florent Ginhoux. Retnla^cre^ mice^27^ were provided by Dr. David Voehringer. Tnfrsf11a^Cre^; Rosa26^LSL-YFP^ and Ms4a3^FlpO^; Rosa26^FSF-tdTomato^ mice^18^ were provided by Dr. Elvira Mass. Experimental and control animals were co-housed, and littermate controls were used as often as possible. Mice were bred and housed in certified housing facilities with an ambient temperature of ∼20-23 °C, a 12/12-hour light/dark cycle and food available ad libitum. Animals were euthanized by cervical dislocation. Animal protocols required for experimentation other than organ collection were authorized by the French Ministry of Higher Education and Research upon approval of the local ethical committee (CIEPAL Azur) at Université Côte d’Azur, and by the Institutional Animal Care and Use Committee (IACUC) at University of Minnesota Medical School.

### *In vivo* treatments

#### Administration of tamoxifen

CCR2^CreERT2^ x R26^TdTomato^ mice were treated with tamoxifen dissolved in corn oil (20 mg/mL) by a single oral gavage (250 μL/mouse). Animals were sacrificed 2 days later and assessed for labeling efficiency in brown adipose tissue by flow cytometry.

CX3CR1^CreERT2^ x R26^TdTomato^ mice were injected intra-peritoneally with 10 mg/mL tamoxifen (200 μL per mouse in 10% EtOH and sunflower oil) daily for 5 days. Labelling efficiency was assessed in brain, blood and brown adipose tissue for each experiment involving this strain. Ms4a3^CreERT2^ x R26^TdTomato^ mice were treated with tamoxifen dissolved in corn oil (20 mg/mL) by oral gavage (100 μL/mouse) daily for 5 days. Labelling efficiency was assessed in blood and brown adipose tissue 3, 10 and 17 days following the last tamoxifen administration.

#### Monocyte depletion with MC-21

Mice were injected intra-peritoneally once a day for 1, 2 or 5 days with 10 μg MC-21 (anti-CCR2) antibody or vehicle (PBS) and were sacrificed 16 h after the last injection for analysis of brown adipose tissue macrophages. Monocyte depletion efficiency was confirmed by blood analysis.

#### Macrophage depletion with AFS98

Mice were injected intra-peritoneally twice (48 hours apart) with 500 μg anti-CD115 blocking antibody (AFS98, BioXCell cat#BE0213) or isotype control (Rat IgG2a, BioXcell cat#BE0089) and were sacrificed 16 h after the last injection for analysis of adipose tissue macrophages.

### In vitro culture

Total bone marrow cells were isolated by flushing bone marrow out of femurs and tibias using PBS. Bone marrow cells were cultured in RPMI medium containing 10% FBS, 2mM L-Glutamine, 50U/mL Penicillin and 50 μg/mL Streptomycin. Growth factors and cytokines were added as indicated in figure legends, at a concentration of 50 ng/mL (M-CSF, GM-CSF) or 10ng/mL (IL-4).

### Flow cytometry

Tissues were harvested after cervical dislocation and washed in PBS. Splenocytes were prepared by gently crushing the spleen on a 70 μm strainer in flow buffer (PBS containing 1% BSA and 2 mM EDTA). Peritoneal cells were obtained by performing lavage with 5 mL PBS. Bone marrow cells were prepared by flushing femurs and tibias with flow buffer. Adipose depots, lungs, liver and brains were harvested, minced with scissors, and then incubated for 45 min with PBS containing 3 mg/mL collagenase A and 100μg/mL DNAse I at 37 °C. For experiments with Tnfrsf11a^Cre^-Ms4a3^Flp^ double fate-mapper mice, adipose tissues were incubated for 30 min with PBS containing 0.5% BSA, 2 mg/mL Collagenase-II and 2.5 mM CaCl_2_. Digested tissues were then homogenized using a 1 mL syringe with a 20G needle and passed through a 70 μm strainer. Brain immune cells were further enriched using a Percoll gradient. Blood was drawn from the submandibular vein and collected in heparinized tubes or Eppendorf tubes containing 15 μl 500 mM EDTA. Red blood cells were lysed using BD Pharm Lyse lysing solution (BdBiosciences cat #555899). Cells were washed with flow buffer and then surface stained. All antibodies were used 1/200. A list of all antibodies used in provided in Supplementary Table 1. Flow cytometry data were acquired using a BD FACS Canto II, BD FACSymphony A5 and a Cytek Aurora cytometer with 5 laser configuration. All analyses were performed using FlowJo software (Tree Star).

### Metabolite analyses

#### Colorimetric quantification of metabolites

Glycerol, TG and NEFA contents were measured from serum. Free glycerol reagent and standard were used according to the manufacturer’s protocol. NEFA-HR2 R1 + R2 FUJIFILM were used according to the manufacturer’s protocol.

#### Serum lipidomic analysis by GCMS-NCI

<u>Materials:</u> Fatty acid calibration standards and deuterated fatty acid internal standards (saturated and unsaturated) were purchased from CDN Isotopes (Cluzeau Info Lab, Sainte Foy La Grande, France) and Cayman (Bertin Pharma, Montigny-le-Bretonneux, France) respectively. Chemicals of the highest grade available were from Sigma Aldrich (Saint-Quentin Fallavier, France). LCMSMS quality grade solvents were purchased from Fischer Scientific (Illkirch, France).

<u>Methods:</u> Serum (25µL) was mixed with 5 µL of a fatty acid internal standard mix, containing 260 ng of myristic acid-d3, 1128 ng of palmitic acid-d3, 840 ng of stearic acid-d3, 720 ng of linoleic acid d4, 10.4 ng of arachidic acid-d3, 432 ng of arachidonic acid-d8, 10.8 ng of behenic acid-d3, 108 ng of DHA-d5, 5.2 ng of Lignoceric-d4, and 4 ng of cerotic acid-d4. To separate non-esterified fatty acids, the spiked serum samples were mixed with 1.2ml of Dole’s reagent (Isopropanol/Hexane/Phosphoric acid 2M 40/10/1 v/v/v). Free fatty acids were then recovered with 1 ml of hexane and 1 ml of distilled water. The organic phase was collected and evaporated under vacuum. Fatty acids were then analyzed as pentafluorobenzyl esters (PFB-FAs esters) by NCI-GCMS as previously described ^39^. Calibration curves were obtained using myristic acid (0.05 - 0.8 µg), palmitic acid (1 - 8 µg), stearic acid (0.5 – 2 µg), linoleic acid (2 - 8 µg), arachidic acid (0.01 – 0.04 µg), arachidonic acid (1 - 4 µg), docosahexaenoic acid (0.4 – 1.6 µg), and docosapentaenoic acid (0.1 – 0.4 µg) extracted by the same method used for plasma. The abundances of serum free fatty acids were determined by negative chemical ionization mode (NCI) GCMS. Linear regression was applied for calculations.

#### Quantitative analysis of brown adipose tissues by targeted mass spectrometry

<u>Materials:</u> Glyceride standards, (17:0)2-d5 DG and (17:0)3 TG were obtained from Avanti Polar Lipids (Coger SAS, Paris, France). High-grade organic solvent modifiers were purchased from Sigma Aldrich (Saint-Quentin Fallavier, France). LC-MSMS quality grade solvents were purchased from Fischer Scientific (Illkirch, France).

<u>Preparation of lipid extracts:</u> Gravimetrically weighed adipose tissues suspended in CHCl_3_:MeOH (2:1) were homogenized with the Omni Bead Ruptor 24 (Omni International. Inc.) coupled to a RT400 savant condensation trap (Thermo Scientific). DG (1µg) and TG (2.5µg) internal standards were added to approximately 0.5mg of adipose tissue homogenates. The spiked homogenates were vortexed, centrifuged, collected into inserts, dried under vacuum, and resuspended in 100µl of CHCl_3_:MeOH (2:1).

Analysis of Glycerides by UHPLC-MS^2^: The homogenates (3 µL) were analyzed by LC-MSMS. For the separation and elution of analytes, a Nucleodur ISIS C18 1,8 µm guard (3 x 4mm) and analytical (3 x 100mm) column (Marechey-Nagel, Hoerdt Cedex, France) maintained at 50°C were used^40^. Briefly, DG and TG were resolved on a 1260 Infinity LC system (Agilent Technologies) with a gradient of mobile phase A (acetonitrile/water/1M ammonium formate (60/39/1 v/v/v) with 0.1% formic acid), and mobile phase B (isopropanol/acetonitrile/1M ammonium formate (90/9/1 v/v/v) with 0.1% formic acid)^41^ at a flow rate of 0.45ml/min set as follows : 1 min hold at 30% B; 30-60% B in 2 min; 60-72% B in 15 min; 72-99% B in 20min;

3 min hold at 99%, 99%-30% ramp-down in 0.1 min and, maintained at 30% B for 3.9 min. Acquisition was realized on a 6460 Triple Quadrupole (Agilent Technologies) equipped with an ESI Jet stream source (temperature 150 °C, nebulizer 15 L/min, sheath gas 11 L/min, sheath gas temperature 150°C, capillary 4500 V and nozzle voltage 1200 V) operated in positive Single Reaction Monitoring (SRM) mode (fragmentor 152 and 220 V, collision energy 29 and 25 V for DG and TG respectively. Ammoniated DG were quantified according to the response observed from the neutral loss of either their sn-1 or sn-2 fatty acid. Ammoniated TG were quantified according to the sum of the responses resulting from the neutral loss of either their sn-1, sn-2, and sn-3 fatty acids. Finally, lipid concentrations were determined by calculating the relative response of each lipiform to its respective internal standard.

### Histology and immunostaining

Tissues were harvested and fixed for 4 h at room temperature or overnight at 4°C in ADAPT-3D fixative ^42^ (4% paraformaldehyde solution with 30% sucrose, pH9). Samples were then washed in 1X PBS. For tissue sections, a paraffin infiltration was performed on dehydrated tissues (Myr Spin Tissue Processor STP 120). Then, tissues were embedded in paraffin (inclusion center EC350-B&PMP). 7 μm sections were performed using a HM340E microtome (Microm Microtech, Francheville France). Paraffin sections were then de-paraffinized and stained using hematoxylin & eosin. For whole mount microscopy, tissues were decolored and stained following the ADAPT-3D method ^42^ and cleared in ethyl cinnamate. Images were acquired on a Nikon CSU-W1 spinning disk microscope.

### Single-cell RNA-seq integration

Single-cell datasets were downloaded from GEO database: GSE177635 ^5^ (brown fat tissue, WT sample GSM5378577); GSE203097 ^23^ (adrenal gland tissue, samples: GSM6153750 and GSM6153751); GSE225665 ^43^ (peritoneal wash, samples: GSM7053956 and GSM7053957). For splenic tissue ^25^, single-cell dataset was downloaded from https://figshare.com/articles/dataset/Bulk_and_single-cell_RNA-seq_data/22232056/1. Cell type annotations were used from original studies (for brown fat and adrenal gland datasets - provided by authors, for peritoneal wash and spleen datasets downloaded together with data from public resources). Datasets were subsetted to maintain only monocyte, macrophage, and dendritic cell populations.

Datasets were integrated using Seurat v4.3.0 tool^44^ using RPCA approach. Samples, coming from different datasets were log normalized, and 3000 variable features were selected using ‘vst’ method. After the selection of integration features, highly variable genes were scaled and variation coming from the percentage of mitochondrial genes and number of counts were regressed out. Integration anchors were found with no k.filter and 250 n.trees parameters. Integrated data were scaled and the final umap reduction was calculated based on the first 25 PCA components.

For DC3 signature enrichment, the top 100 differentially expressed genes were calculated using the FindMarkers function, with the following parameters: logfc.threshold = 0.01, min.pct = 0.01, only.pos = T. Enrichment plot was created using plotCoregulationProfileReduction function from fgsea package v1.27.1^45^. Scatter plots were visualized using ggplot2 v3.4.2 package^46^. Shapes of cluster borders were calculated by first estimating 2d kernel density using MASS package v7.3-58.4^47^, followed by creating a grid graph with igraph package v1.4.3^48^ and calculating α-shape of clusters using alphahull v2.5 package^49^. Visualization of gene expression for Gallerand et al., 2021 dataset in Figures S1, and S4 was generated using scNavigator (Artyomov lab, https://artyomovlab.wustl.edu/scn/).

### Statistical Analysis

All data are represented in mean ± SEM. Statistical analysis was performed with GraphPad Prism 10 as indicated in each figure legend. Presence of statistically significant differences between conditions are indicated as follows: ns p>0.05 ; * p<0.05 ; ** p<0.01 ; *** p<0.001 ; **** p<0.0001.

## References

1. Mass, E., Nimmerjahn, F., Kierdorf, K., and Schlitzer, A. (2023). Tissue-specific macrophages: how they develop and choreograph tissue biology. Nat. Rev. Immunol. 23, 563–579. 10.1038/s41577-023-00848-y.

2. Ginhoux, F., and Guilliams, M. (2016). Tissue-Resident Macrophage Ontogeny and Homeostasis. Immunity 44, 439–449. 10.1016/j.immuni.2016.02.024.

3. Liu, Z., Gu, Y., Chakarov, S., Bleriot, C., Kwok, I., Chen, X., Shin, A., Huang, W., Dress, R.J., Dutertre, C.A., et al. (2019). Fate Mapping via Ms4a3-Expression History Traces Monocyte-Derived Cells. Cell 178, 1509–1525 e1519. 10.1016/j.cell.2019.08.009.

4. Félix, I., Ollikainen, J., Haque, A., Karhula, J., Salmi, M., Jokela, H., and Rantakari, P. (2025). Embryonic macrophages in brown adipose tissue are spatially associated with developing nerves. bioRxiv, 2025.2001.2016.632922. 10.1101/2025.01.16.632922.

5. Gallerand, A., Stunault, M.I., Merlin, J., Luehmann, H.P., Sultan, D.H., Firulyova, M.M., Magnone, V., Khedher, N., Jalil, A., Dolfi, B., et al. (2021). Brown adipose tissue monocytes support tissue expansion. Nat Commun 12, 5255. 10.1038/s41467-021-25616-1.

6. Rosen, E.D., and Spiegelman, B.M. (2014). What we talk about when we talk about fat. Cell 156, 20–44. 10.1016/j.cell.2013.12.012.

7. Nicholls, D.G. (2017). The hunt for the molecular mechanism of brown fat thermogenesis. Biochimie 134, 9–18. 10.1016/j.biochi.2016.09.003.

8. Wolf, Y., Boura-Halfon, S., Cortese, N., Haimon, Z., Sar Shalom, H., Kuperman, Y., Kalchenko, V., Brandis, A., David, E., Segal-Hayoun, Y., et al. (2017). Brown-adipose-tissue macrophages control tissue innervation and homeostatic energy expenditure. Nat Immunol 18, 665–674. 10.1038/ni.3746.

9. Cox, N., Crozet, L., Holtman, I.R., Loyher, P.L., Lazarov, T., White, J.B., Mass, E., Stanley, E.R., Elemento, O., Glass, C.K., and Geissmann, F. (2021). Diet-regulated production of PDGFcc by macrophages controls energy storage. Science 373. 10.1126/science.abe9383.

10. Ingersoll, M.A., Spanbroek, R., Lottaz, C., Gautier, E.L., Frankenberger, M., Hoffmann, R., Lang, R., Haniffa, M., Collin, M., Tacke, F., et al. (2010). Comparison of gene expression profiles between human and mouse monocyte subsets. Blood 115, e10–19. 10.1182/blood-2009-07-235028.

11. Sunderkotter, C., Nikolic, T., Dillon, M.J., Van Rooijen, N., Stehling, M., Drevets, D.A., and Leenen, P.J. (2004). Subpopulations of mouse blood monocytes differ in maturation stage and inflammatory response. J Immunol 172, 4410–4417. 10.4049/jimmunol.172.7.4410.

12. Carlin, L.M., Stamatiades, E.G., Auffray, C., Hanna, R.N., Glover, L., Vizcay-Barrena, G., Hedrick, C.C., Cook, H.T., Diebold, S., and Geissmann, F. (2013). Nr4a1-dependent Ly6C(low) monocytes monitor endothelial cells and orchestrate their disposal. Cell 153, 362–375. 10.1016/j.cell.2013.03.010.

13. Auffray, C., Fogg, D., Garfa, M., Elain, G., Join-Lambert, O., Kayal, S., Sarnacki, S., Cumano, A., Lauvau, G., and Geissmann, F. (2007). Monitoring of blood vessels and tissues by a population of monocytes with patrolling behavior. Science 317, 666–670. 10.1126/science.1142883.

14. Hanna, R.N., Cekic, C., Sag, D., Tacke, R., Thomas, G.D., Nowyhed, H., Herrley, E., Rasquinha, N., McArdle, S., Wu, R., et al. (2015). Patrolling monocytes control tumor metastasis to the lung. Science 350, 985–990. 10.1126/science.aac9407.

15. Trzebanski, S., Kim, J.S., Larossi, N., Raanan, A., Kancheva, D., Bastos, J., Haddad, M., Solomon, A., Sivan, E., Aizik, D., et al. (2024). Classical monocyte ontogeny dictates their functions and fates as tissue macrophages. Immunity. 10.1016/j.immuni.2024.04.019.

16. Menezes, S., Melandri, D., Anselmi, G., Perchet, T., Loschko, J., Dubrot, J., Patel, R., Gautier, E.L., Hugues, S., Longhi, M.P., et al. (2016). The Heterogeneity of Ly6C(hi) Monocytes Controls Their Differentiation into iNOS(+) Macrophages or Monocyte-Derived Dendritic Cells. Immunity 45, 1205–1218. 10.1016/j.immuni.2016.12.001.

17. Jung, S., Aliberti, J., Graemmel, P., Sunshine, M.J., Kreutzberg, G.W., Sher, A., and Littman, D.R. (2000). Analysis of fractalkine receptor CX(3)CR1 function by targeted deletion and green fluorescent protein reporter gene insertion. Mol Cell Biol 20, 4106–4114. 10.1128/MCB.20.11.4106-4114.2000.

18. Huang, H., Balzer, N.R., Seep, L., Splichalova, I., Blank-Stein, N., Viola, M.F., Franco Taveras, E., Acil, K., Fink, D., Petrovic, F., et al. (2025). Kupffer cell programming by maternal obesity triggers fatty liver disease. Nature. 10.1038/s41586-025-09190-w.

19. Mass, E., Ballesteros, I., Farlik, M., Halbritter, F., Gunther, P., Crozet, L., Jacome-Galarza, C.E., Handler, K., Klughammer, J., Kobayashi, Y., et al. (2016). Specification of tissue-resident macrophages during organogenesis. Science 353. 10.1126/science.aaf4238.

20. Mack, M., Cihak, J., Simonis, C., Luckow, B., Proudfoot, A.E., Plachy, J., Bruhl, H., Frink, M., Anders, H.J., Vielhauer, V., et al. (2001). Expression and characterization of the chemokine receptors CCR2 and CCR5 in mice. J. Immunol. 166, 4697–4704. 10.4049/jimmunol.166.7.4697.

21. Schraml, B.U., van Blijswijk, J., Zelenay, S., Whitney, P.G., Filby, A., Acton, S.E., Rogers, N.C., Moncaut, N., Carvajal, J.J., and Reis e Sousa, C. (2013). Genetic tracing via DNGR-1 expression history defines dendritic cells as a hematopoietic lineage. Cell 154, 843–858. 10.1016/j.cell.2013.07.014.

22. Kim, K.W., Williams, J.W., Wang, Y.T., Ivanov, S., Gilfillan, S., Colonna, M., Virgin, H.W., Gautier, E.L., and Randolph, G.J. (2016). MHC II+ resident peritoneal and pleural macrophages rely on IRF4 for development from circulating monocytes. J Exp Med 213, 1951–1959. 10.1084/jem.20160486.

23. Dolfi, B., Gallerand, A., Firulyova, M.M., Xu, Y., Merlin, J., Dumont, A., Castiglione, A., Vaillant, N., Quemener, S., Gerke, H., et al. (2022). Unravelling the sex-specific diversity and functions of adrenal gland macrophages. Cell Rep. 39, 110949. 10.1016/j.celrep.2022.110949.

24. Han, J., Gallerand, A., Erlich, E.C., Helmink, B.A., Mair, I., Li, X., Eckhouse, S.R., Dimou, F.M., Shakhsheer, B.A., Phelps, H.M., et al. (2024). Human serous cavity macrophages and dendritic cells possess counterparts in the mouse with a distinct distribution between species. Nat. Immunol. 25, 155–165. 10.1038/s41590-023-01688-7.

25. Liu, Z., Wang, H., Li, Z., Dress, R.J., Zhu, Y., Zhang, S., De Feo, D., Kong, W.T., Cai, P., Shin, A., et al. (2023). Dendritic cell type 3 arises from Ly6C(+) monocyte-dendritic cell progenitors. Immunity 56, 1761–1777 e1766. 10.1016/j.immuni.2023.07.001.

26. Tabula Muris, C. (2020). A single-cell transcriptomic atlas characterizes ageing tissues in the mouse. Nature 583, 590–595. 10.1038/s41586-020-2496-1.

27. Krljanac, B., Schubart, C., Naumann, R., Wirtz, S., Culemann, S., Kronke, G., and Voehringer, D. (2019). RELMalpha-expressing macrophages protect against fatal lung damage and reduce parasite burden during helminth infection. Sci Immunol 4. 10.1126/sciimmunol.aau3814.

28. Westermann, F., Tuzlak, S., Kreiner, V., Bejarano, D., Bijnen, M., Cecconi, V., van Hove, H., Wang, H., Litscher, G., Ignacio, A., et al. (2024). Unveiling a unique macrophage population in exocrine glands sustained by ILC2-derived GM-CSF. bioRxiv, 2024.2010.2030.620897. 10.1101/2024.10.30.620897.

29. Komuczki, J., Tuzlak, S., Friebel, E., Hartwig, T., Spath, S., Rosenstiel, P., Waisman, A., Opitz, L., Oukka, M., Schreiner, B., et al. (2019). Fate-Mapping of GM-CSF Expression Identifies a Discrete Subset of Inflammation-Driving T Helper Cells Regulated by Cytokines IL-23 and IL-1beta. Immunity 50, 1289–1304 e1286. 10.1016/j.immuni.2019.04.006.

30. Helft, J., Bottcher, J., Chakravarty, P., Zelenay, S., Huotari, J., Schraml, B.U., Goubau, D., and Reis e Sousa, C. (2015). GM-CSF Mouse Bone Marrow Cultures Comprise a Heterogeneous Population of CD11c(+)MHCII(+) Macrophages and Dendritic Cells. Immunity 42, 1197–1211. 10.1016/j.immuni.2015.05.018.

31. Briseno, C.G., Haldar, M., Kretzer, N.M., Wu, X., Theisen, D.J., Kc, W., Durai, V., Grajales-Reyes, G.E., Iwata, A., Bagadia, P., et al. (2016). Distinct Transcriptional Programs Control Cross-Priming in Classical and Monocyte-Derived Dendritic Cells. Cell Rep. 15, 2462–2474. 10.1016/j.celrep.2016.05.025.

32. Schneider, C., Nobs, S.P., Kurrer, M., Rehrauer, H., Thiele, C., and Kopf, M. (2014). Induction of the nuclear receptor PPAR-gamma by the cytokine GM-CSF is critical for the differentiation of fetal monocytes into alveolar macrophages. Nat. Immunol. 15, 1026–1037. 10.1038/ni.3005.

33. Gautier, E.L., Shay, T., Miller, J., Greter, M., Jakubzick, C., Ivanov, S., Helft, J., Chow, A., Elpek, K.G., Gordonov, S., et al. (2012). Gene-expression profiles and transcriptional regulatory pathways that underlie the identity and diversity of mouse tissue macrophages. Nat Immunol 13, 1118–1128. 10.1038/ni.2419.

34. Miller, J.C., Brown, B.D., Shay, T., Gautier, E.L., Jojic, V., Cohain, A., Pandey, G., Leboeuf, M., Elpek, K.G., Helft, J., et al. (2012). Deciphering the transcriptional network of the dendritic cell lineage. Nat. Immunol. 13, 888–899. 10.1038/ni.2370.

35. Randolph, G.J., Ivanov, S., Zinselmeyer, B.H., and Scallan, J.P. (2017). The Lymphatic System: Integral Roles in Immunity. Annu. Rev. Immunol. 35, 31–52. 10.1146/annurev-immunol-041015-055354.

36. Serbina, N.V., and Pamer, E.G. (2006). Monocyte emigration from bone marrow during bacterial infection requires signals mediated by chemokine receptor CCR2. Nat. Immunol. 7, 311–317. 10.1038/ni1309.

37. Vanneste, D., Bai, Q., Hasan, S., Peng, W., Pirottin, D., Schyns, J., Marechal, P., Ruscitti, C., Meunier, M., Liu, Z., et al. (2023). MafB-restricted local monocyte proliferation precedes lung interstitial macrophage differentiation. Nat. Immunol. 24, 827–840. 10.1038/s41590-023-01468-3.

38. Croxford, A.L., Lanzinger, M., Hartmann, F.J., Schreiner, B., Mair, F., Pelczar, P., Clausen, B.E., Jung, S., Greter, M., and Becher, B. (2015). The Cytokine GM-CSF Drives the Inflammatory Signature of CCR2+ Monocytes and Licenses Autoimmunity. Immunity 43, 502–514. 10.1016/j.immuni.2015.08.010.

39. Nguyen, M., Bourredjem, A., Piroth, L., Bouhemad, B., Jalil, A., Pallot, G., Le Guern, N., Thomas, C., Pilot, T., Bergas, V., et al. (2021). High plasma concentration of non-esterified polyunsaturated fatty acids is a specific feature of severe COVID-19 pneumonia. Sci. Rep. 11, 10824. 10.1038/s41598-021-90362-9.

40. Tamba Sompila, A.W., Heron, S., Hmida, D., and Tchapla, A. (2017). Fast non-aqueous reversed-phase liquid chromatography separation of triacylglycerol regioisomers with isocratic mobile phase. Application to different oils and fats. J. Chromatogr. B Analyt. Technol. Biomed. Life Sci. 1041*-*1042, 151–157. 10.1016/j.jchromb.2016.12.030.

41. Cajka, T., Hricko, J., Rudl Kulhava, L., Paucova, M., Novakova, M., and Kuda, O. (2023). Optimization of Mobile Phase Modifiers for Fast LC-MS-Based Untargeted Metabolomics and Lipidomics. Int. J. Mol. Sci. 24. 10.3390/ijms24031987.

42. Daniel D. Lee, D.L.D., Leon C.D. Smyth, et al. (2025). ADAPT-3D: Accelerated Deep Adaptable Processing of Tissue for 3-Dimensional Fluorescence Tissue Imaging for Research and Clinical Settings. PREPRINT (Version 1) available at Research Square. [10.21203/rs.3.rs-6109657/v1].

43. Li, X., Mara, A.B., Musial, S.C., Kolling, F.W., Gibbings, S.L., Gerebtsov, N., and Jakubzick, C.V. (2024). Coordinated chemokine expression defines macrophage subsets across tissues. Nat. Immunol. 25, 1110–1122. 10.1038/s41590-024-01826-9.

44. Hao, Y., Hao, S., Andersen-Nissen, E., Mauck, W.M., 3rd, Zheng, S., Butler, A., Lee, M.J., Wilk, A.J., Darby, C., Zager, M., et al. (2021). Integrated analysis of multimodal single-cell data. Cell 184, 3573–3587 e3529. 10.1016/j.cell.2021.04.048.

45. Korotkevich, G., Sukhov, V., Budin, N., Shpak, B., Artyomov, M.N., and Sergushichev, A. (2021). Fast gene set enrichment analysis. bioRxiv, 060012. 10.1101/060012.

46. Wickham, H. (2016). ggplot2: Elegant Graphics for Data Analysis. Springer-Verlag New York 978-3-319-24277-4.

47. Venables WN, R.B. (2002). Modern Applied Statistics with S, Fourth edition. Springer, New York ISBN 0-387-95457-0.

48. Csárdi G, N.T., Traag V, Horvát S, Zanini F, Noom D, Müller K (2024). igraph: Network Analysis and Visualization in R. R package version 2.0.3. doi:10.5281/zenodo.7682609.

49. Pateiro-López, B., & Rodríguez-Casal, A (2010). Generalizing the Convex Hull of a Sample: The R Package alphahull. Journal of Statistical Software 34(5), 1–28.

