## Supplementary Data for "CD226^+^ macrophages arise from MDP-derived monocytes and regulate lipid metabolism"

### Supplementary Figure 1

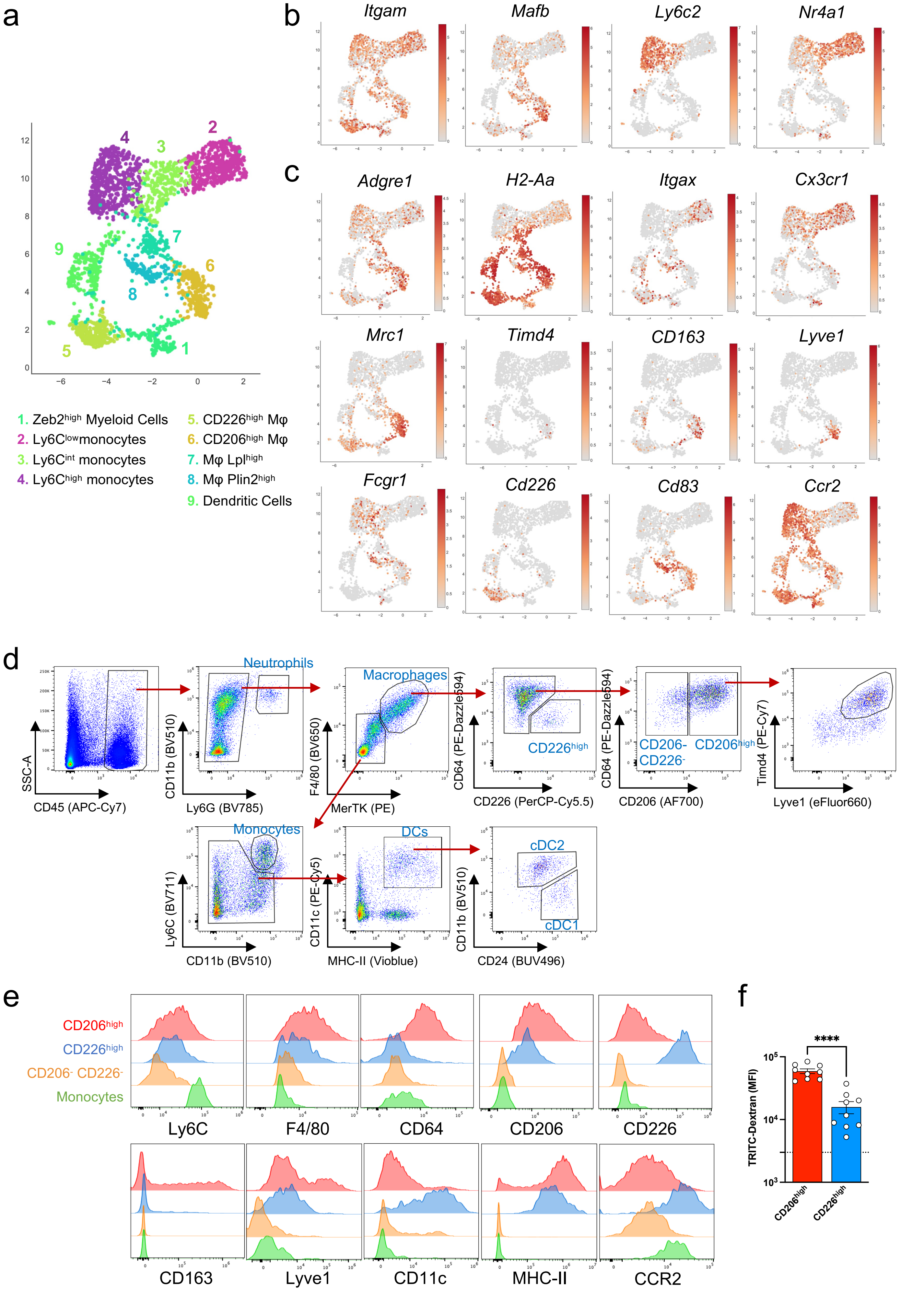

**Supplementary Figure 1. Characterization of brown adipose tissue macrophage diversity.** (**a-c**) Analysis of brown adipose tissue myeloid cells using single cell RNA-sequencing. (**d**) Gating strategy used to identify immune cell populations in adipose tissue. (**e**) Histograms representing expression of indicated markers by BAT monocytes and macrophages, analyzed by flow cytometry. (**f**) Quantification of TRITC fluorescence intensity in BAT $CD206^{high}$ and $CD226^{high}$ macrophages following intravenous injection of TRITC-Dextran. Data are presented as mean values +/- SEM. This figure refers to main figure 1.

### Supplementary Figure 2

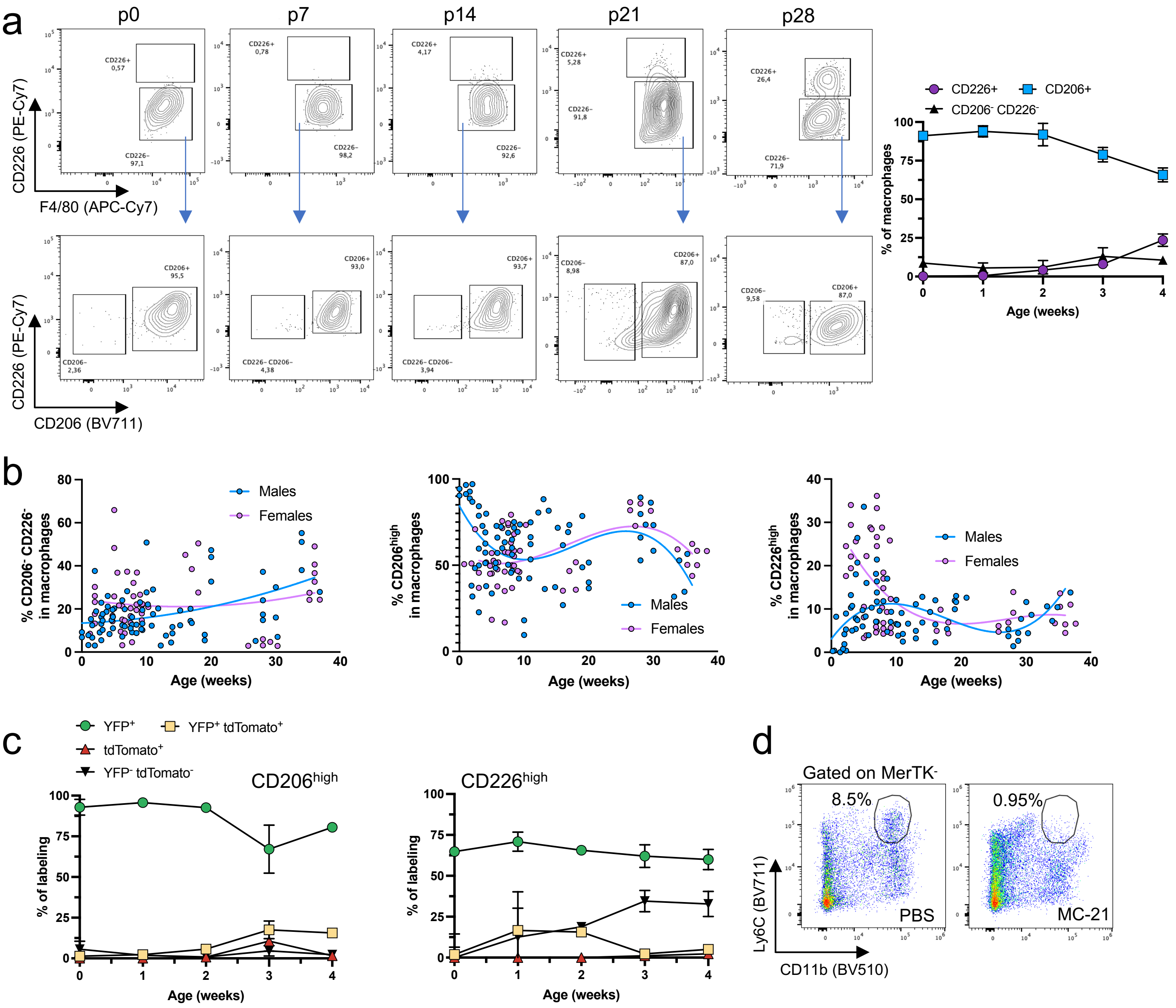


**Supplementary Figure 2. Characterization of brown adipose tissue macrophages across age and sex.** (**a**) Time-course flow cytometry analysis of BAT CD226$^{high}$ and CD206$^{high}$ macrophage content in the first weeks after birth. (**b**) Proportions of CD206$^{high}$, CD226$^{high}$ and CD206$^{-}$ CD226$^{-}$ macrophages subsets in BAT depending on sex and age. (**c**) Flow cytometry analysis of tdTomato and YFP expression in BAT CD226$^{high}$ and CD206$^{high}$ macrophages from Ms4a3$^{Flp}$; R26$^{FSF\text{-}tdTomato}$; Tnfrsf11a$^{Cre}$; R26$^{LSL\text{-}YFP}$ mice.(**d**) Validation of monocyte depletion in BAT from wild-type mice following MC-21 administration. Data are presented as mean values +/- SEM. This figure refers to main figure 1.

### Supplementary Figure 3

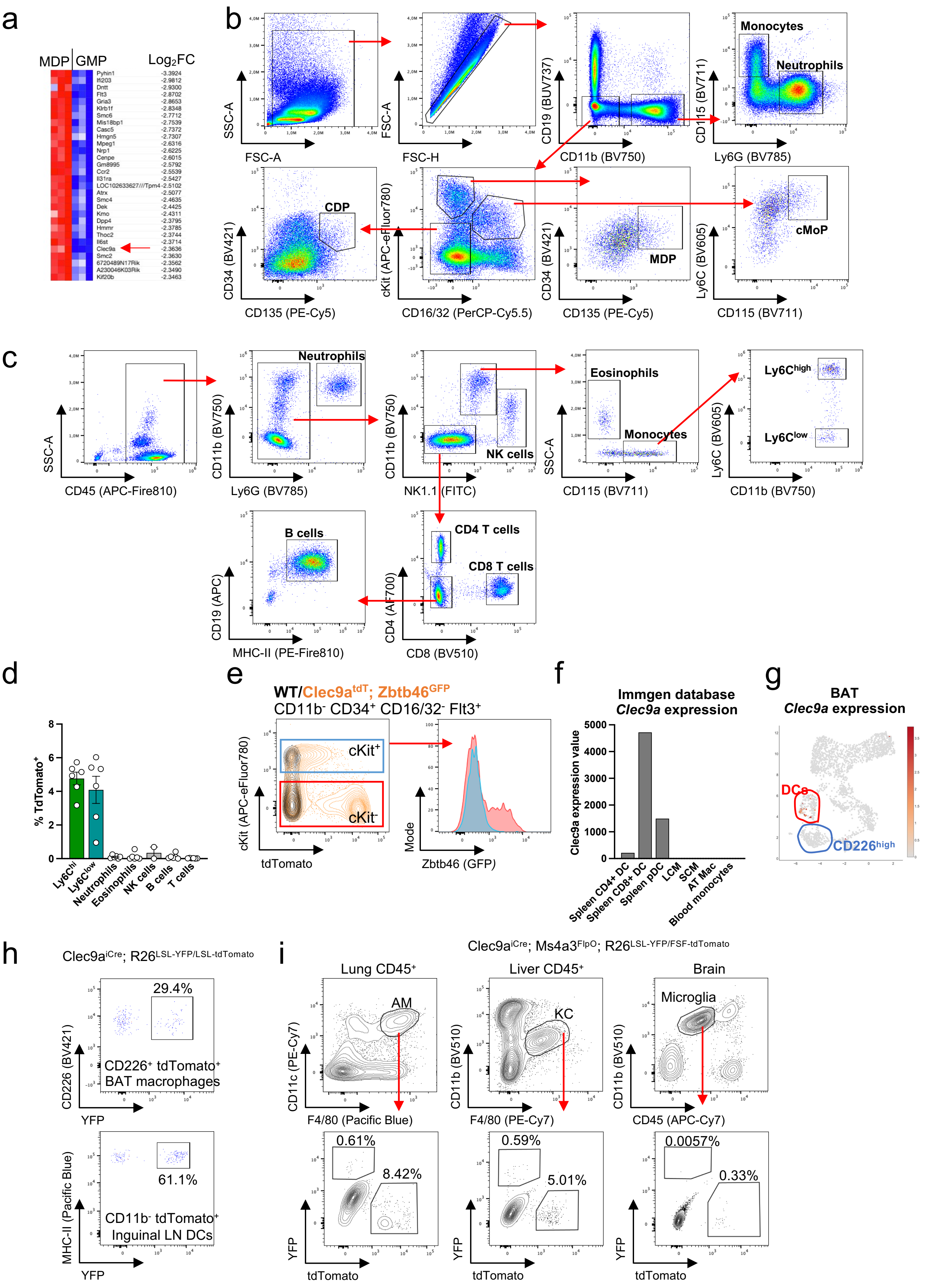

**Supplementary Figure 3. Past Clec9a expression identifies MDP-derived monocytes and macrophages.**
(**a**) Differential gene expression analysis between bone marrow GMPs and MDPs. Data from the Immgen consortium. (**b**) Gating strategy used to identify bone marrow precursors. (**c**) Gating strategy used to identify blood immune cells. (**d**) TdTomato labeling of blood immune cells in $Clec9a^{iCre}$; $R26^{tdTomato}$ mice. (**e**) ) Flow cytometry analysis of bone marrow $CD16/32^{-}$ $Flt3^{+}$ progenitors from $Clec9a^{iCre}$; $R26^{tdTomato}$; $Zbtb46^{GFP}$ mice. (**f**) Expression of *Clec9a* by splenic dendritic cells, peritoneal cavity macrophages, adipose tissue macrophages and blood monocytes. Data from Immgen.org. (**g**) UMAP representing *Clec9a* expression by BAT myeloid cells. Data are presented as mean values +/- SEM. This figure refers to main figure 2.

### Supplementary Figure 4

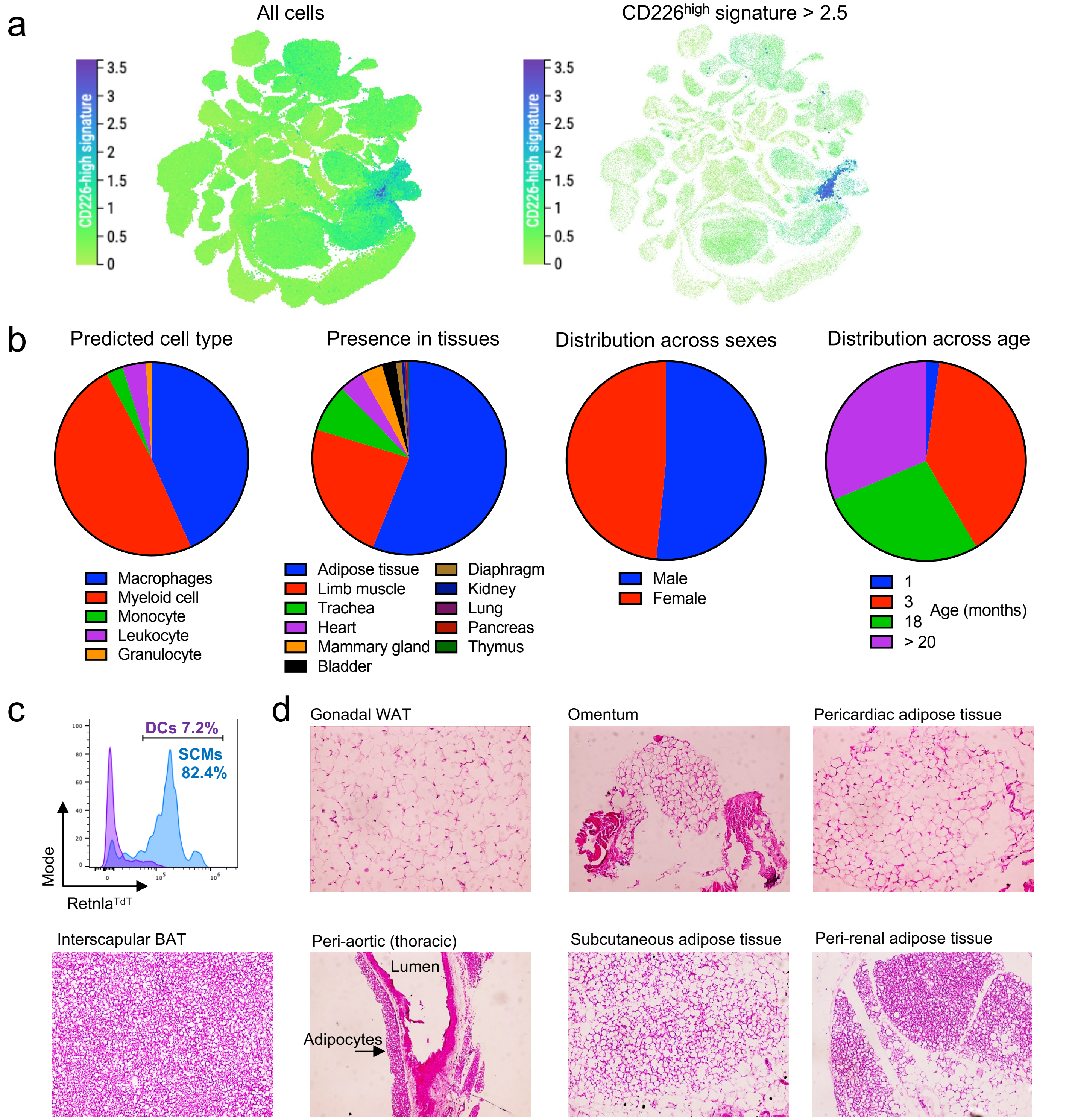


**Supplementary Figure 4. Characterization of brown adipose tissue macrophages across age and sex.** (**a**) Expression of the "CD226$^{high}$ signature" consisting of the top 50 genes differentially expressed in CD226$^{high}$ macrophages, in the Tabula Muris Senis dataset. (**b**) Characterization of CD226$^{high}$ macrophages identified in the Tabula Muris Senis dataset. (**c**) Expression of tdTomato in peritoneal SCMs and dendritic cells (DCs) from Retnla$^{Cre}$; R26$^{tdTomato}$ mice. (**d**) Hematoxylin and eosin staining of interscapular BAT, peri-aortic brown adipose tissue, subcutaneous adipose tissue, peri-renal adipose tissue, peri-cardiac adipose tissue, omentum and gonadal white adipose tissue. Data are presented as mean values +/- SEM. This figure refers to main figure 3.

#### Supplementary Figure 5

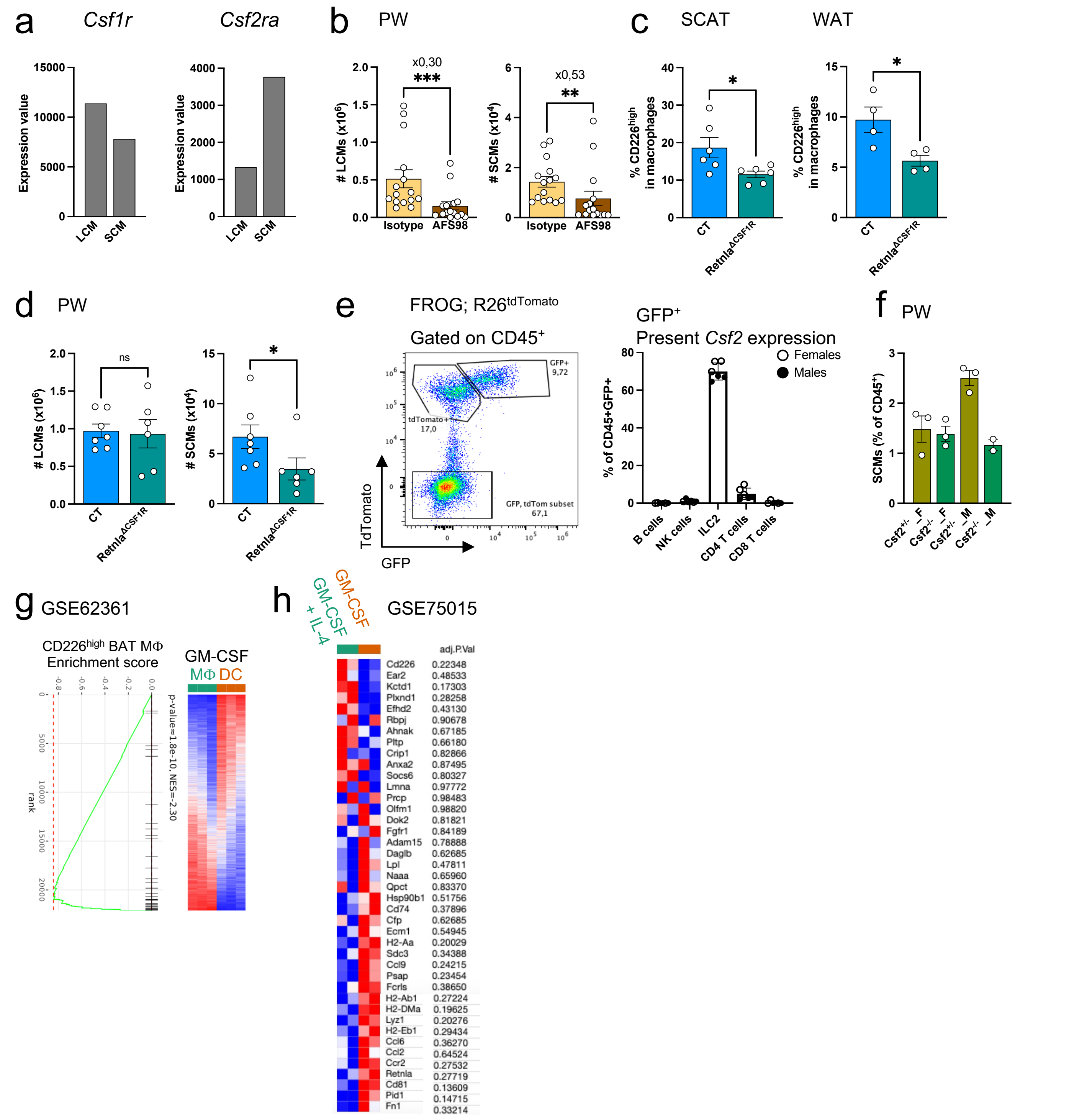

**Supplementary Figure 5. CSF1R and GM-CSF regulate CD226$^{high}$ macrophage numbers.**
(**a**) Expression of *Csf1r* and *Csf2ra* by peritoneal cavity macrophages. Data from Immgen.org. (**b**) Quantification of peritoneal LCMs and SCMs following isotype or AFS98 treatment. (**c-d**) Quantification of SCAT, WAT (c) and PW (d) macrophage subsets in control (CT) and Retnla$^{\Delta CSF1R}$ mice. (**e**) Quantification of GM-CSF-expressing GFP$^{+}$ cells in adipose tissue from FROG; R26$^{tdTomato}$ mice (**f**) Quantification of peritoneal SCMs from male and female Csf2$^{+/-}$ and Csf2$^{-/-}$ mice. (**g**) GSEA analysis, comparing the signature of BAT CD226$^{high}$ macrophages to macrophages and DCs obtained by culturing bone marrow with GM-CSF (GSE62361). (**h**) Differential expression of BAT CD226$^{high}$ macrophage signature genes between "MoDCs" differentiated in the presence of GM-CSF +/- IL-4 (GSE75015). Data are presented as mean values +/- SEM. This figure refers to main figure 4.

### Supplementary Figure 6

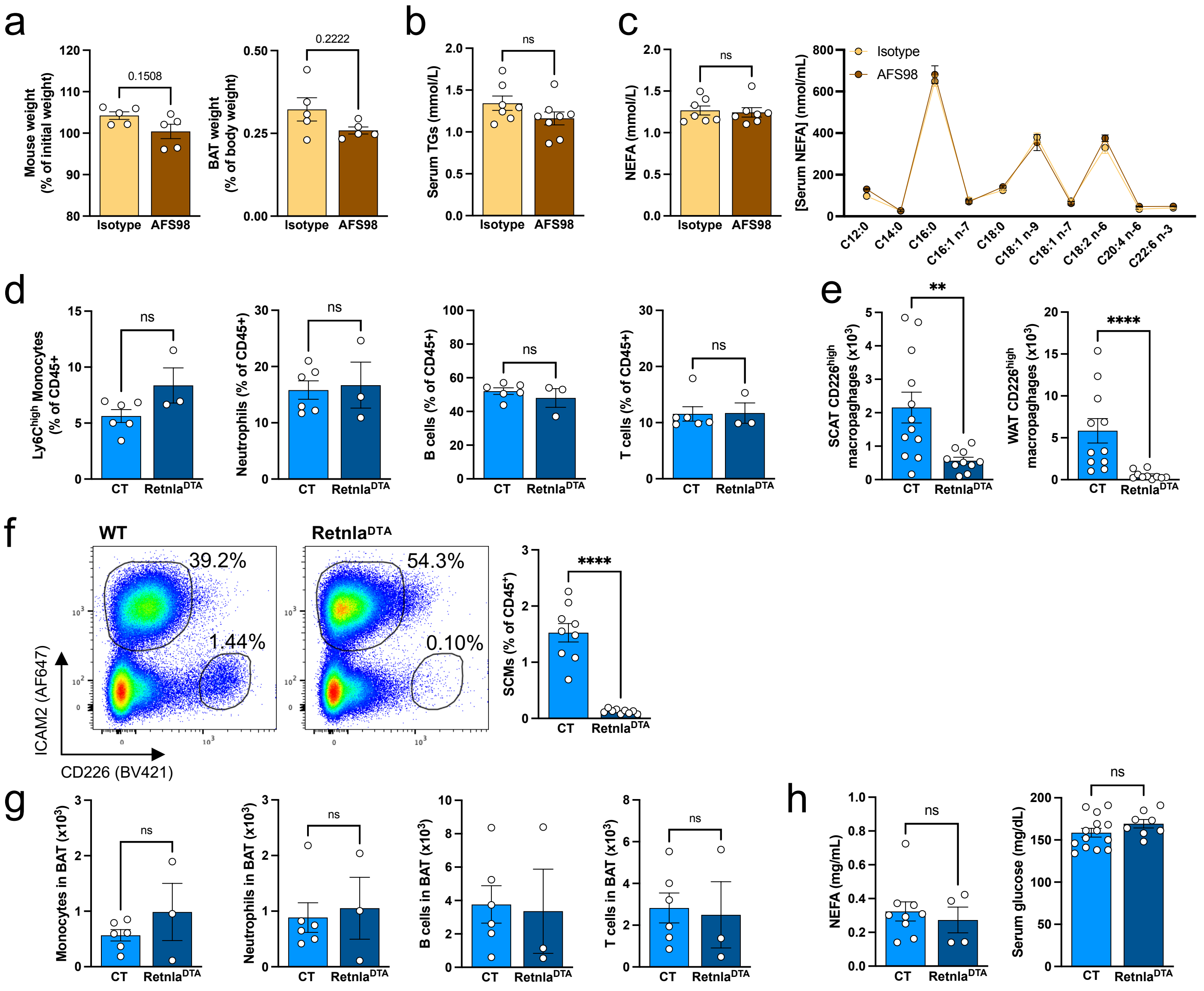


**Supplementary Figure 6. Phenotyping Retnla$^{DTA}$ mice.**

(**a**) Body weight and BAT weight in wild-type mice treated with isotype control or AFS98. (**b, c**) Serum TG (b) and NEFA (c) levels in wild-type mice treated with isotype control or AFS98. (**d**) Quantification of blood immune cells in control and Retnla$^{DTA}$ mice. (**e**) Quantification of macrophage subsets in SCAT and WAT from control and Retnla$^{DTA}$ mice. (**f**) Quantification of SCMs in peritoneal wash from control and Retnla$^{DTA}$ mice. (**g**) Quantification of monocytes, neutrophils and B and T lymphocytes in BAT from control and Retnla$^{DTA}$ mice. (**h**) Serum NEFA and glucose levels in control and Retnla$^{DTA}$ mice. Data are presented as mean values +/- SEM. This figure refers to main figure 5.

**Supplementary Table 1**

**Key Resources Table**

| Reagent or resource | Source | Identifier |
|---|---|---|
| **Antibodies** | | |
| α-mouse CD115 PE (clone AFS98) | Biolegend | Cat# 135506 |
| α-mouse CD115 BV711 (clone AFS98) | Biolegend | Cat# 135515 |
| α-mouse/human CD11b Brilliant Violet 510 (clone M1/70) | Biolegend | Cat# 101263 |
| α-mouse/human CD11b PE-Cy5 (clone M1/70) | Biolegend | Cat# 101210 |
| α-mouse/human CD11b BV750 (clone M1/70) | Biolegend | Cat# 101267 |
| α-mouse Treml4 PE (clone 16E5) | Biolegend | Cat# 143304 |
| α-mouse Gr1 PerCP-Cy5.5 (clone RB6-8C5) | BD Biosciences | Cat# 552093 |
| α-mouse Gr1 FITC (clone RB6-8C5) | Biolegend | Cat# 108406 |
| α-mouse CD16/32 PerCP-Cy5.5 (clone 93) | Biolegend | Cat# 156624 |
| α-mouse CD34 BV421 (clone SA376A4) | Biolegend | Cat# 152208 |
| α-mouse Ly6C BV421 (clone HK1.4) | Biolegend | Cat# 128032 |
| α-mouse Ly6C BV711 (clone HK1.4) | Biolegend | Cat# 128037 |
| α-mouse Ly6C BV605 (clone HK1.4) | Biolegend | Cat# 128036 |
| α-mouse Ly6G BV785 (clone 1A8) | Biolegend | Cat# 127645 |
| α-mouse ICAM2 AF647 (clone 3C4 (MIC2/4)) | Biolegend | Cat# 105612 |
| α-mouse CD9 APC-Fire750 (clone MZ3) | Biolegend | Cat# 124814 |
| α-mouse CD3e BV605 (clone 145-2C11) | Biolegend | Cat# 100351 |
| α-mouse CD3e APC (clone 145-2C11) | Biolegend | Cat# 100312 |
| α-mouse CD4 AF700 (clone RM4-4) | Biolegend | Cat# 116022 |
| α-mouse CD8 AF647 (clone 53-6.7) | Biolegend | Cat# 100724 |
| α-mouse CD45 BV570 (clone 30-F11) | Biolegend | Cat# 103136 |
| α-mouse F4/80 PE-Cy7 (clone BM8) | Biolegend | Cat# 123114 |
| α-mouse F4/80 AF488 (clone BM8) | Biolegend | Cat# 123120 |
| α-mouse CD45 APC-Cy7 (clone 30-F11) | BD Biosciences | Cat# 557659 |
| α-mouse CD64 Brillant Violet 421 (clone X54-5/7.1) | Biolegend | Cat# 139309 |
| α-mouse CD64 PE-Cy7 (clone X54-5/7.1) | Biolegend | Cat# 139314 |
| α-mouse CD14 PE (clone M14-23) | Biolegend | Cat# 150106 |
| α-mouse MerTK PE (clone 2B10C42) | Biolegend | Cat# 151506 |
| α-mouse CD11c PE-Cy5 (clone N418) | Biolegend | Cat# 117316 |
| α-mouse MHC-II (IA/IE) PB (clone M5/114.15.2) | Biolegend | Cat# 107620 |
| α-mouse MHC-II (IA/IE) VioBlue (clone M5/114.15.2) | Miltenyi Biotec | Cat# 130-123-278 |
| α-mouse MHC-II (IA/IE) PE-Fire810 (clone M5/114.15.2) | Biolegend | Cat# 107667 |
| α-mouse CD226 BV421 (clone TX42.1) | Biolegend | Cat# 133615 |
| α-mouse TCRβ PB (clone H57-597) | Biolegend | Cat# 109226 |
| α-mouse NK1.1 APC (clone PK136) | Biolegend | Cat# 108720 |
| α-mouse Ter119 APC (clone TER-119) | Biolegend | Cat# 116212 |

| | | |
|---|---|---|
| α-mouse B220 APC (clone RA3-6B2) | BD Biosciences | Cat# 561226 |
| α-mouse CD19 BUV737 (clone 1D3) | BD Biosciences | Cat# 612781 |
| α-mouse CD150 PE-Cy7 (clone TC15-12F12.2) | Biolegend | Cat# 115914 |
| α-mouse Sca1 PB (clone D7) | Biolegend | Cat# 108120 |
| α-mouse Sca1 PE-Cy7 (clone D7) | Biolegend | Cat# 108114 |
| α-mouse c-Kit APC-Cy7 (clone ACK2) | eBioscience | Cat# 47-1172-82 |
| α-mouse CD48 AF488 (clone HM48-1) | Biolegend | Cat# 103414 |
| α-mouse CXCR4 APC (clone 2B11) | eBioscience | Cat# 51-9991-80 |
| α-mouse CCR2 PE (clone REA538) | Miltenyi Biotec | Cat# 130-117-548 |
| α-mouse CCR2 APC-Fire750 (clone SA203G11) | Biolegend | Cat# 150630 |
| α-mouse CD226 PerCP-Cy5.5 (clone TX42.1) | Biolegend | Cat# 133624 |
| α-mouse CD226 BV421 (clone TX42.1) | Biolegend | Cat# 133615 |
| α-mouse CD206 AF647 (clone C068C2) | Biolegend | Cat# 141712 |
| α-mouse CD206 APC (clone C068C2) | Biolegend | Cat# 141708 |
| α-mouse CD135 PE-Cy5 (clone A2F10) | Biolegend | Cat# 135312 |
| α-mouse CD11b APC (clone M1/70) | Biolegend | Cat# 101218 |

**Reagents**

| | | |
|---|---|---|
| DAPI | Sigma | Cat# D9542 |
| PFA 4% | VWR International | Cat# 9713.1000 |
| Bovine serum Albumin (BSA) | Sigma | Cat# A7030 |
| Collagenase A | Sigma | Cat# 11088793001 |
| DNAse I | Sigma | Cat# 10104159001 |
| Liberase | Roche | Cat# 5401054001 |
| ImmunoHistoMount | Sigma | Cat# I1161 |
| RBC lysing buffer | BD Biosciences | Cat# 555899 |
| Free glycerol reagent | Sigma | Cat# F6428 |
| DreamTaq Green PCR Master Mix (2X) | Thermo Scientific | Cat# K1081 |
| Brilliant Stain Buffer Plus | BD Biosciences | Cat# 566385 |
| EDTA | Sigma | Cat# 324504-500ML |
| Live/Dead fixable viability dye | Thermofisher | Cat #L34955 |

**Kits**

| | | |
|---|---|---|
| NEFA-HR2 R1 + R2 FUJFILM | WAKO | Cat# W1W270-77000 |
| Glucose dosage Kit | BioSentec | Cat# 075 |
| Triglyceride dosage Kit | DiaSys | Cat# 157109910021 |

**Accessories**

| | | |
|---|---|---|
| StepOne | Applied Biosystem | N/A |
| Thermo Cycler SimpliAmp | Applied Biosystem | N/A |
| Nanodrop | OZYME | N/A |
| Veterinary hematology analyzer | Exigo | H400 |
| Aurora 5 laser configuration | Cytek | N/A |
| BD FACS Canto II | BD Biosciences | N/A |

**Softwares**

| | | |
|---|---|---|
| Prism10 | GraphPad | N/A |
| FlowJo | Tree Star | N/A |
| StepOne Software v.2.2.2 | Applied Biosystem | N/A |
| Fiji | https://imagej.net/software/fiji/ | N/A |
